# Convergent genomic and molecular features predict risk of metachronous metastasis in clear cell renal cell carcinoma

**DOI:** 10.1101/2025.01.07.630976

**Authors:** Marjan M. Naeini, Mengyuan Pang, Neha Rohatgi, Sinem Kadioglu, Umesh Ghoshdastider, Renzo G. DiNatale, Roy Mano, A Ari Hakimi, Anders Jacobsen Skanderup

## Abstract

The molecular features determining the risk of metachronous metastases in clear cell renal cell carcinoma (ccRCC) are poorly defined. Using a systematic tumor transcriptome deconvolution approach, we investigated the genomic and transcriptomic profiles of 192 ccRCC primary tumors with extended clinical follow-up to identify cancer and stromal cell molecular features associated with metastatic risk. At the genomic level, we identified a significantly higher frequency of copy number loss at 1p31-36 in primary tumors that later progressed with metastases. Tumor transcriptome deconvolution identified significant down-regulation of epithelial cell polarity, including *PATJ* (1p31), and fatty acid metabolism, including *CYP4A11* (1p33), in cancer cells of tumors that developed metastatic progression. We developed and benchmarked a compact 5-feature predictive model (5G) that demonstrated improved accuracy over existing ccRCC gene signatures in the prediction of metachronous metastasis risk. Overall, our study highlights convergent genomic and transcriptomic alterations in chromosome 1p, driving dysregulation of epithelial cell polarity and fatty acid metabolism, as putative novel risk factors of metachronous metastasis in ccRCC.

## Introduction

Metastatic clear cell renal carcinoma (ccRCC) has a 5-year survival rate of ∼10% and is generally considered incurable (Wu et al. 2023). Metastatic spread of ccRCC is often diagnosed during pre- operative assessment (i.e. synchronous spread). However, around one-third of ccRCC patients with localized disease eventually relapse with metastatic progression following curative surgery (Tran and Ornstein 2022). The third prespecified interim analysis of the KEYNOTE-564 trial demonstrated that adjuvant pembrolizumab significantly and clinically meaningfully improved overall survival compared to placebo in participants with ccRCC at high risk of recurrence after surgery (Choueiri et al. 2024). The ability to identify ccRCC patients with an increased risk of developing metachronous metastasis could lead to improved treatment strategies, such as active surveillance and additional adjuvant therapy using pembrolizumab (Hakimi and Voss 2018, Dudani et al. 2019, Choueiri et al. 2024).

Previous studies have established the genomic and molecular landscape of ccRCC (Creighton et al. 2013, Sato et al. 2013, Turajlic et al. 2018) and demonstrated widespread metabolic reprogramming (Hakimi et al. 2016), immune infiltration signatures (Şenbabaoğlu et al. 2016), as well as recurrent alterations of the PI3K/AKT/mTOR pathway (Creighton et al. 2013, Sato et al. 2013, Scelo et al. 2014). Recurrent copy number alterations include loss of chromosome arms 1p, 3p, 4q, 6q, 8p, 9p and gains of 1q, 2q, 5q, 7q, 8q, 12p, and 20q (Beroukhim et al. 2009, Creighton et al. 2013, Scelo et al. 2014, Turajlic et al. 2018). In the metastatic setting, genomic loss of 9p and 14q have been reported as clonally selected and expanded driver events within metastatic lesions (Turajlic et al. 2018). Patients with metastatic disease and low body mass index (BMI) tend to have a poor prognosis, and studies have suggested a role for dysregulated fatty acid metabolism in the metastatic progression of ccRCC (Albiges et al. 2016).

Efforts to decipher the transcriptomic features of metastatic ccRCC tumors have led to the development of putative prognostic biomarkers, including the ClearCode34 risk predictor for localized ccRCC (Brooks et al. 2014), a 16-gene recurrence score (Rini et al. 2015), and an RNA-based 31-gene cell cycle progression (CCP) signature (Morgan et al. 2018). Furthermore, metastatic progression is often accompanied by marked changes in the tumor microenvironment (TME) (Quail and Joyce 2013). A recent study identified intratumoral myeloid inflammation as a key driver of metastasis in ccRCC (Rappold et al. 2022). Moreover, Alchahin et al. investigated single cell gene expression data from tumors diagnosed with synchronous metastasis and proposed biomarkers associated with synchronous metastasis (Alchahin et al. 2022). These studies underline the influence of the TME on the metastatic progression of ccRCC. However, our understanding of cancer and stroma cell features rendering patients with localized tumors at risk of progressing with metachronous metastasis remains nascent.

Here we sought to identify cancer and stromal cell molecular characteristics of primary ccRCC tumors associated with the development of metachronous metastasis. We conducted an integrative genome and transcriptome analysis of 192 primary tumors with prolonged clinical follow-up (up to 8 years) to accurately distinguish indolent tumors from those that subsequently developed metastasis. We comprehensively investigated potential genomic features associated with the development of metachronous metastasis. We then performed a deep comparative analysis of the tumor transcriptomes in these patient subsets, comparing both inferred immune cell infiltrates as well as deconvoluted cancer and stromal-cell tumor transcriptomes. Finally, we developed and benchmarked a metachronous metastasis risk model based on the top predictive gene expression features in the dataset. Overall, our study provides a systematic analysis of the molecular features associated with metachronous metastasis in ccRCC, identifying candidate biomarkers for improved risk stratification of primary tumors at risk of metastatic progression.

## Results

### Clinical characteristics of cohorts

We performed extended clinical follow-up for metastatic events (median follow-up time of 23.2 months) in 192 ccRCC primary tumors profiled by TCGA (Creighton et al. 2013) (**Figure 1** and **Supplementary** Figure 1). We classified stage III primary tumors without subsequent metastasis during the entire follow-up as “indolent” (IN; n=80) (**Figure 1a-c**). Non-metastatic primary tumors that developed metastasis more than 100 days following diagnosis were classified as “metachronous metastatic” (MM; n=44) (**Figure 1a-c)**. Finally, stage IV primary tumors or tumors developing metastasis within 100 days following diagnosis were classified as “synchronous metastatic” (SM; n=68) (**Figure 1a-c**). Consistent with previous reports (Albiges et al. 2016), patients with SM tumors had significantly lower body mass index (BMI) as compared to other tumors (Kruskal-Wallis, *P*=0.0014) (**Supplementary** Figure 2), while no significant difference was found between IN and MM tumors (Wilcoxon rank-sum*, P*=0.3). Similarly, SM tumors were larger (Kruskal-Wallis, *P*=6.4e-04), with no difference between MM and IN tumors (Wilcoxon rank-sum, *P*=0.22). Overall, these observations are consistent with previously reported features of metastatic ccRCC and support the notion that MM tumors are indistinguishable from IN tumors at the macroscopic level. To further characterize and validate the molecular findings of our analysis, we used an external cohort comprising 69 non-metastatic and 32 metastatic tumor samples with gene expression data (Sato et al. 2013) (**Supplementary** Figure 3).

**Figure 1:**
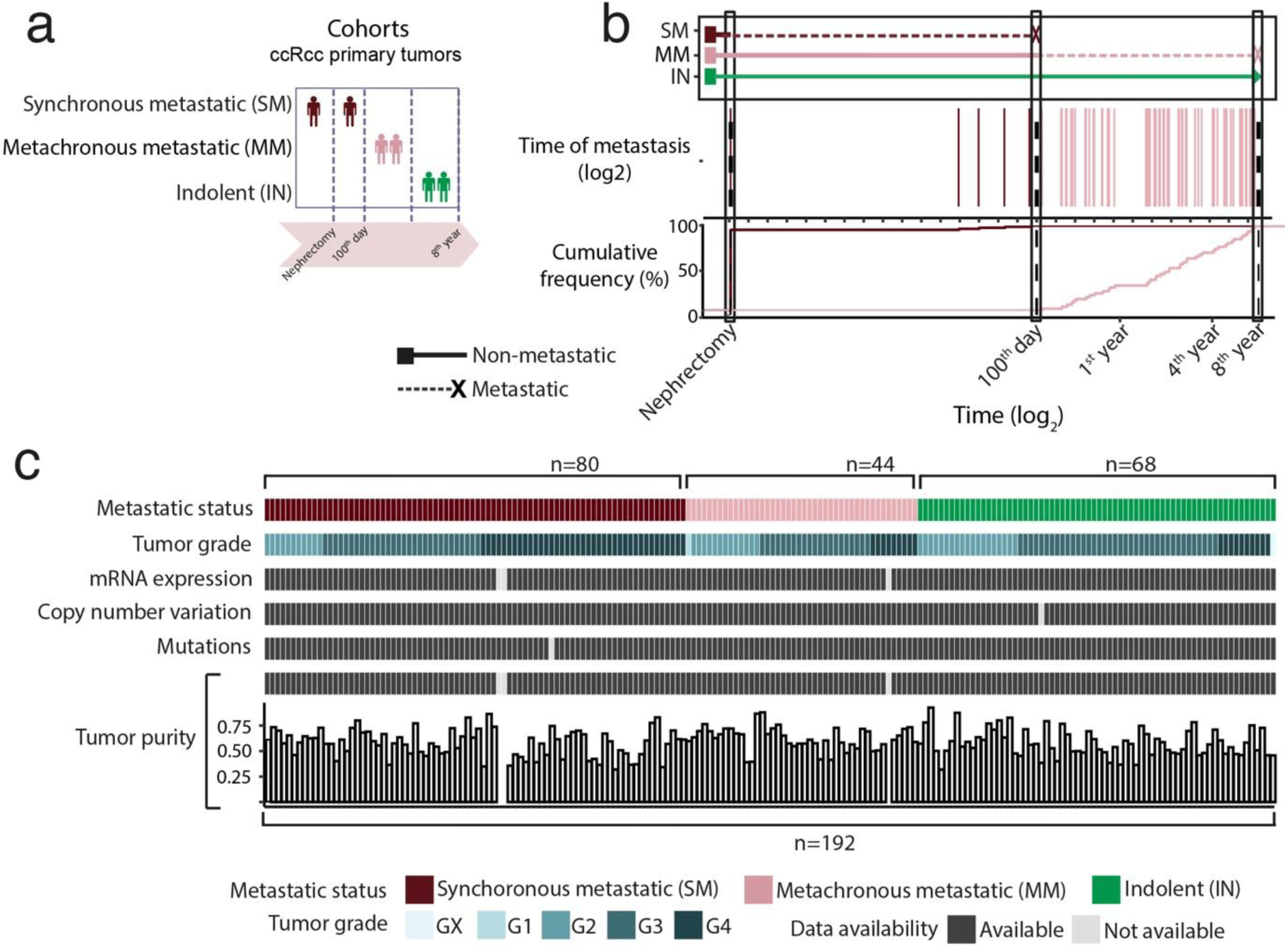
Overview of ccRCC cohort. **a)** Schematic presentation of ccRCC cohorts**. b**) Tumor samples grouped by metastatic progression status. Synchronous metastatic (SM) primary tumors were metastatic at presentation or <100 days following nephrectomy. Metachronous metastatic (MM) primary tumors developed delayed metastasis, >100 days following nephrectomy. Indolent (IN) primary tumors had no reported metastasis during the clinical follow-up time. **c**) Available genomic and molecular data across the ccRCC cohort and metastatic progression status.

### Chromosome 1p33 loss is associated with metachronous metastasis

We first examined the genomic profiles in the three tumor subsets. Somatic mutation frequencies were not significantly associated with synchronous or metachronous metastasis in our cohort (**Figure 2a**). Mutations in *BAP1* and *SETD2* have previously been linked to worse cancer-specific survival (Hakimi et al., CCR, 2013). Indeed, our analysis also highlighted the increased mutation frequency of both genes in metastatic patients (Figure 2a). *FREM1*, an extracellular matrix protein, showed enrichment of frameshift deletion and missense mutations in MM tumors (n=4) as compared to IN tumors (n=0) (**Figure 2a** and **Supplementary** Figure 4). However, this difference was not significant after multiple testing correction (Fisher’s exact test, *P* = 0.022, *q*-value=1).

**Figure 2:**
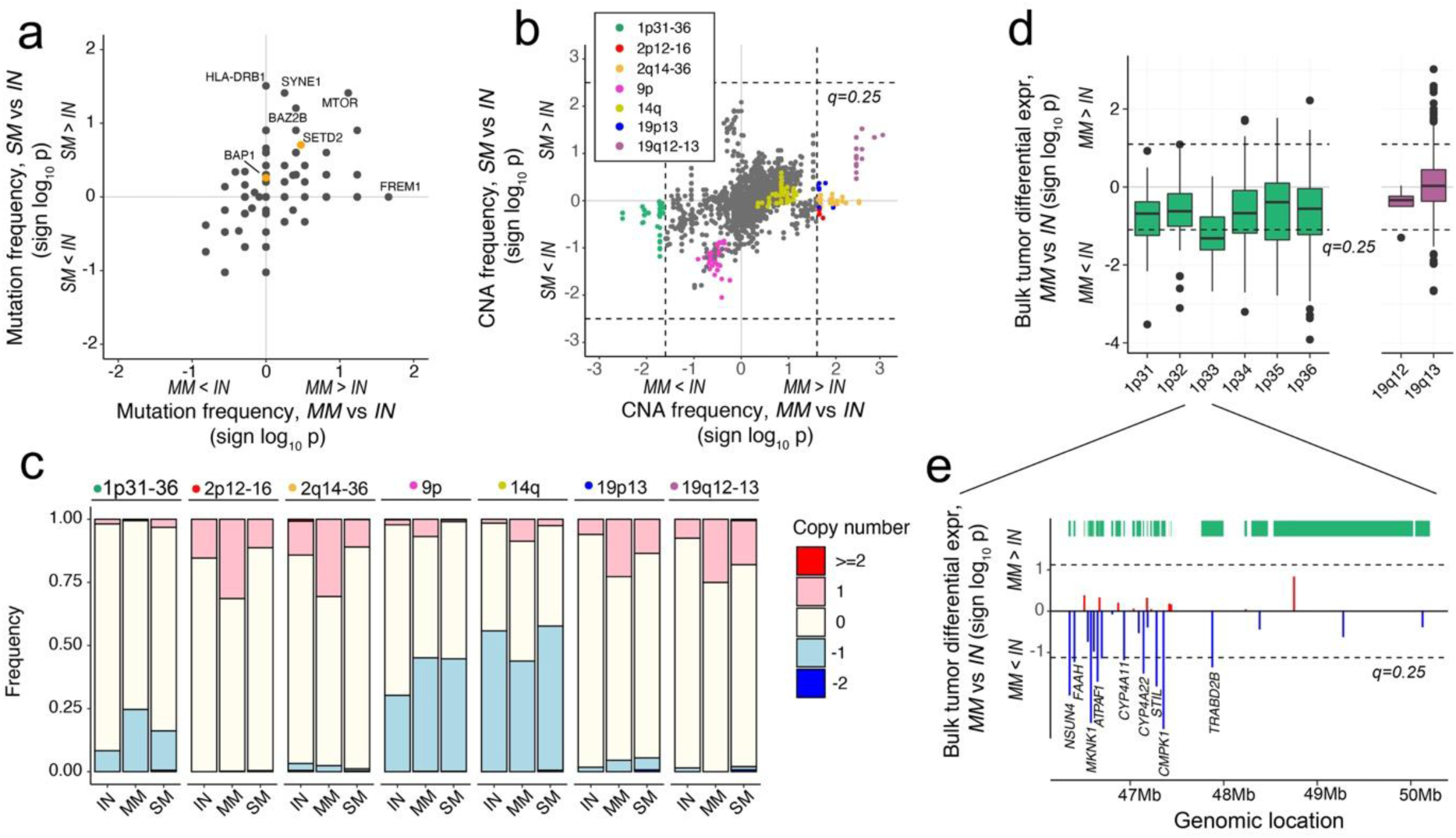
**Association of metastasis with mutations and copy number alterations**. **a**) Comparison of gene mutation frequencies in metastatic (SM and MM) versus indolent (IN) tumors (two-tailed Fisher’s exact test). No genes were significantly enriched in metastatic tumors after correcting for multiple testing. Known driver genes, *BAP1* and *SETD2*, are highlighted in orange color. **b**) Comparison of copy number alteration frequencies in metastatic (SM and MM) versus indolent (IN) tumors (two-tailed Wilcoxon-rank-sum test). Loci with significant (*q*<0.25) differences between MM and IN cohorts are highlighted. Additionally, regions (9p and 14q) previously associated with distant metastasis are highlighted. **c**) Copy number alteration frequencies of significant and highlighted regions across IN, MM and SM cohorts. **d**) Comparison of bulk tumor expression of genes located in the 1p31-36 (CNA loss) and 19q12-13 (CNA gain) regions in MM and IN cohorts (two-tailed t-test). **e**) Bulk tumour differential expression (sign log_10_ p, t-test) between MM and IN samples for individual genes located on 1p33. Significantly dysregulated genes are labelled (*q* <0.25). Box plots in d display the median values with the interquartile range (lower and upper hinge) and ±1.5-fold the interquartile range from the first and third quartile (lower and upper whiskers). IN, indolent cohort; MM, metachronous metastasis cohort; SM, synchronous metastasis cohort; CNA, copy number alteration.

Next, we examined the frequency of chromosomal copy number alterations (CNA) in the three groups. We identified CNAs at five regions associated with MM tumors (Wilcoxon rank-sum, *q*-value<0.2) (**Figure 2b**). These alterations comprised copy number gain at 19q12-13, 19p13, 2q14-36 and 2p12- 16, and copy number loss across 1p31-36 (**Figure 2c** and **Supplementary** Figure 5a). We identified a strong positive correlation between copy number alterations and gene expression levels in 1p31-36, which was not observed in the other four regions (**Supplementary** Figure 5b). Similarly, genes located at the center of this region (1p33) showed lower expression in MM compared to IN tumors (Wilcoxon rank-sum*, q*-value<0.2) (**Figure 2d** and **Supplementary** Figure 6). A closer inspection of 1p33 revealed multiple altered genes (*NSUN4*, *FAAH*, *MKNK1*, *ATPAF1*, *CYP4A11*, *CYP4A22*, *STIL*, *CMPK1*, *TRABD2B*) with genomic loss and significantly down-regulated expression in MM compared to IN tumors (Wilcoxon rank-sum*, q*-value<0.25) (**Figure 2e**). While recurrent 1p36 copy number loss has been reported in multiple ccRCC cohorts (Creighton et al. 2013; Sato et al. 2013; Turajlic et al. 2018), a previous report did not identify 1p36 loss as a selected and clonally expanded driver event in distant metastatic lesions (Turajlic et al. 2018). Taken together, these results suggest a relationship between chromosomal instability and metastatic seeding potential in localized ccRCC lesions, notably involving copy number loss and concomitant down-regulation of gene expression at or near 1p33-36.

### Depletion of regulatory T cells and M1 macrophages in MM tumors

To explore the role of TME immune cell infiltration as a risk factor for metastatic progression, we analyzed the immune cell type composition using the tumor RNA-seq profiles. We used three distinct methods for deconvolution of immune cell subsets from bulk tumor transcriptomic data: xCell (Aran et al. 2017), Consensus^TME^ (Jiménez-Sánchez et al. 2019) and Cibersort (Newman et al. 2015). xCell identified strong depletion of T regulatory cells (Tregs) (*P*=2.99e-09) (**Figure 3a**) and M1 macrophages (*P*=6.6e-0.3) in MM as compared to IN tumors. Similarly, Consensus^TME^ confirmed the depletion of M1 macrophages and Tregs (*P*=0.1 and *P*=0.022) (**Figure 3a**). Although not significant, depletion of these two cell types was also identified by Cibersort (*P*=0.40 and *P*=0.30) (**Supplementary** Figure 7). We then compared the expression of individual Treg and M1 macrophages marker genes in MM and IN tumors (**Supplementary** Figures 8 and 9). Among Treg gene markers, *MCM9*, *TULP4*, *STAM* and *IL10RA* showed significant down-regulation in MM tumors (*P*<0.01) (**Figure 3b**). Similarly, we identified significant down-regulation of multiple M1 macrophages marker genes (*NDUFS2*, *FKBP15* and *SLC31A1)* in MM tumors (*P*<1e-03) (**Figure 3c**). Further investigation of these marker genes in SM tumors demonstrated consistent changes in gene expression for MM and SM tumors, but often with stronger expression changes in MM samples (**Figure 3b-c**).

**Figure 3:**
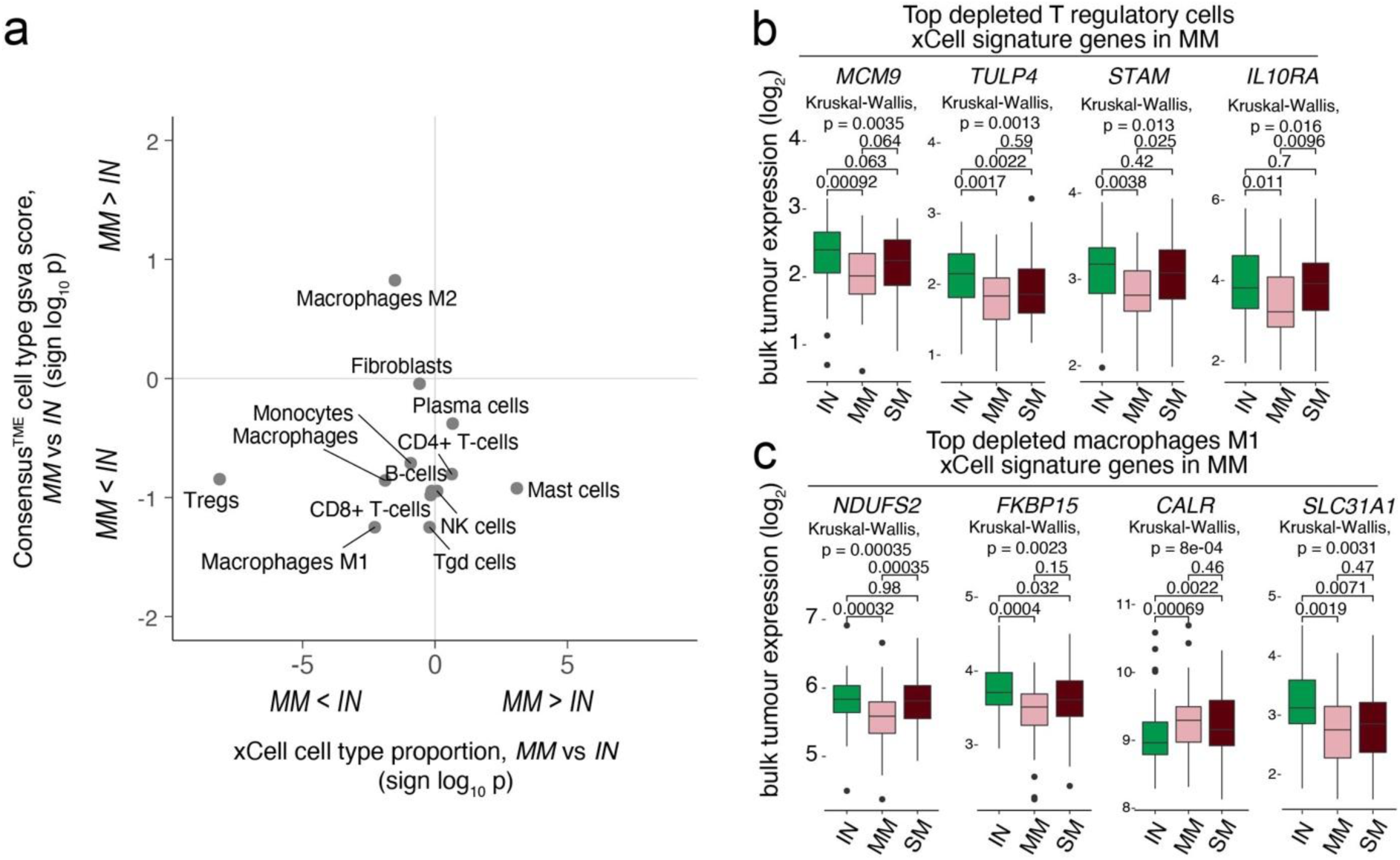
**Immune cell type infiltration**. **a**) Comparison of immune cell type proportions in MM and IN tumors (Wilcoxon rank sum test) using xCell and ConsensusTME. **b**) Top downregulated (Kruskal-Wallis test) T-regulatory cell signature genes in MM cohort. **c**) Top downregulated (Kruskal-Wallis test) Macrophage M1 signature genes in MM cohort. Box plots in b and c refer to the median values with the interquartile range (lower and upper hinge) and ±1.5-fold the interquartile range from the first and third quartile (lower and upper whiskers). Tregs, T regulatory cells; IN, indolent cohort; MM, metachronous metastasis cohort; SM, synchronous metastasis cohort.

### Dysregulation of epithelial cell polarity associated with risk of metastatic progression

We conducted a systematic and unbiased analysis of gene expression differences in the three tumor subsets. A principal component analysis of bulk tumor transcriptomic data using the top 3000 variable genes could not stratify IN, MM and SM patients (**Supplementary** Figure 10a-d). We then performed a comparative analysis of potential differences in the transcriptome profiles using a previously published tumor transcriptome deconvolution approach (Ghoshdastider et al. 2021), which uses tumor purity to infer expression profiles of cancer and stromal cells inside the tumors (see Methods, **Figure 4a**). Tumor purity was estimated using the genomic and transcriptomic profiles for each sample (see Methods). The samples showed a mean tumor purity of 49% (range 24%-82%) with no significant differences between metastatic groups (*P*=0.067).

**Figure 4:**
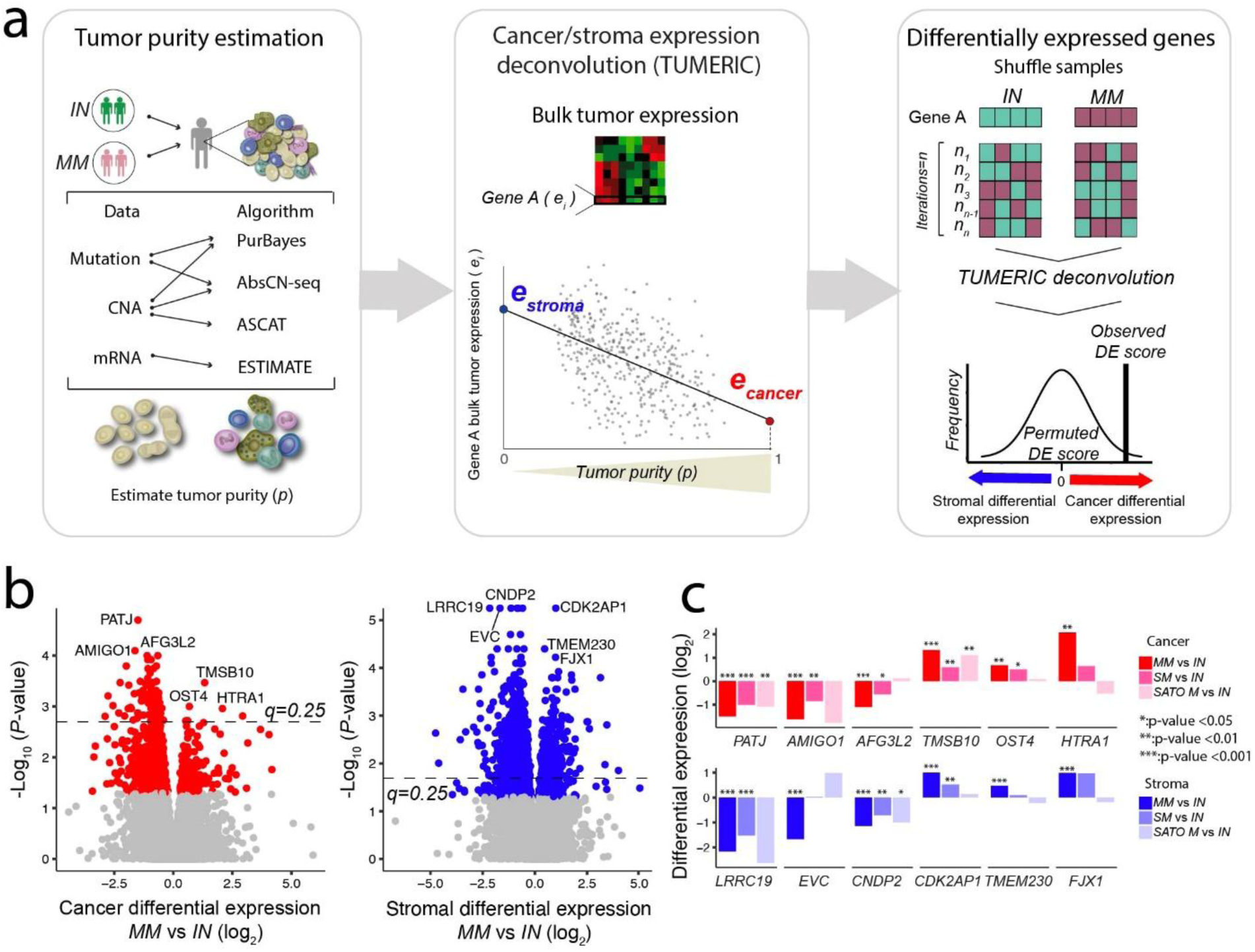
Cancer vs. stroma transcriptome deconvolution. **a**) Tumor purity of each sample is jointly estimated from DNA and RNA data using a consensus approach. Bulk tumor gene expression is then deconvoluted into cancer and stroma cell components. Genes with differential cancer or stromal cell expression across groups (IN and MM) are identified using a sample permutation approach. **b**) Cancer and stromal differential expression comparing MM vs. IN cohorts. Blue and red colors represent significant genes (*P*<0.05). **c**) The top differentially expressed genes are compared across groups, including an independent cohort of metastatic ccRCC tumors (SATO M, Sato et al. 2013). Differential expression relative to the IN group is displayed; ***: p<0.001; **: p <0.01; *: p<0.05. IN, indolent cohort; MM, metachronous metastasis cohort; SM, synchronous metastasis cohort; SATO M, SATO et al. metastatic cohort.

We then compared the deconvoluted cancer cell gene expression profiles across the metastatic groups to identify putative gene expression signatures associated with the risk of metastatic progression. Relative to IN tumors, MM cancer cells demonstrated 87 down-regulated and 6 up- regulated genes (*q* < 0.25, **Figure 4b** and **Supplementary Table 2**). *PATJ*, a tight junction and Crumbs cell polarity component, was the most significant down-regulated gene in MM tumors (*P*=2e-05) (**Figure 4b**). Expression of *PATJ* was also significantly down-regulated in cancer cells of SM samples (*P=*5.8e-04) (**Figure 4c**), suggesting a role for *PATJ* in metastatic progression. To further explore the robustness of this association, we compared bulk tumor expression of *PATJ* in an independent cohort of metastatic and indolent ccRCC samples (Sato et al. 2013). This analysis also confirmed significant down-regulation of *PATJ* in metastatic tumors (*P*=2.8e-03) (**Figure 4c**). Intriguingly, *PATJ* is also located on 1p31, displaying frequent copy number loss in MM tumors, indicating a convergence of genomic and transcriptomic alterations underlying risk of metastatic progression.

*TMSB10* was the most significantly up-regulated gene in cancer cells of MM tumors (*P*=3.4e-04), with expression also up-regulated in SM (*P*=8.1e-03) and metastatic tumors from Sato et al. (*P*=1.4e-03) (**Figure 4c**). *TMSB10* encodes for a member of the beta thymosin protein family which regulate cytoskeleton organization. Previous reports have also suggested a role for this protein in regulation of epithelial cell polarity in breast and ccRCC tumors (Zhang et al. 2017, Xiao et al. 2019). Observing that multiple of the top dysregulated genes in MM tumors were involved in regulation of cell polarity, tight junction formation, and actin assembly, we examined the expression profiles of other known components of these processes (Zhang et al. 2017, Xiao et al. 2019). This analysis revealed significant down-regulation of additional factors such as *AMOLT2*, *CGN* and *TJP2* (**Supplementary** Figure 11a-c). Overall, these results highlight a putative role for dysregulation of tight junction and apicobasal cell polarity in cancer cells as a risk factor for the development of metachronous metastasis.

### Gene expression signatures of tumor stromal cells associated with metastatic risk

We next compared the deconvoluted stromal cell (comprising all non-malignant cells in the tumors) gene expression profiles across metastatic groups to identify tumor stroma gene expression signatures associated with the risk of metastatic progression. We identified 719 down-regulated and 240 up-regulated genes in the stroma of MM as compared to IN tumors (*q*<0.25, **Supplementary Table 3**)*. LRRC19*, a pathogen-recognition receptor and inducer of pro-inflammatory cytokines, was strongly down-regulated in MM tumors (*P*=1.0e-05). The expression of *LRRC19* was also down-regulated in stromal cells of SM samples (*P*=1.0e-05) (**Figure 4c**) and showed down-regulation in an independent cohort of metastatic tumors (*P*=8.9e-02) (**Figure 4c**). We also observed stromal down-regulation of *CNDP2*, encoding a nonspecific dipeptidase involved in histidine metabolism, in MM tumors (*P*=1.0e- 05) as well as the two cohorts of metastatic tumors (*P*=1.7e-03; *P*=3.4e-02) (**Figure 4c**). Further analysis of public single-cell RNA-seq data from normal kidney tissue (Thul et al. 2017) showed that both *LRRC19* and *CNDP2* had the highest expression in proximal tubular cells (**Supplementary** Figure 12). Furthermore, other top differentially expressed genes in the stroma (**Figure 4c**) had the highest expression in either proximal tubular (*LRRC19*, *CNDP2*, *FJX1*), distal tubular (*TMEM230*), or collecting duct cells (*EVC*, *CDK2AP1*) (**Supplementary** Figure 12). Taken together, these results indicate that many of the inferred stromal gene expression differences could be related to compositional differences of healthy kidney tissue cell types in MM and IN tumors.

### Dysregulation of fatty acid metabolism associated with metachronous metastasis

To further explore the biological features associated with the MM tumor subset, we performed pathway-based enrichment analyses of genes with differential expression in cancer cells of MM and IN tumors (**Figure 4b**). We observed significant down-regulation of genes associated with metabolic pathways (*P*=1e-04, Fisher’s exact test) and fatty acid degradation (*P*=6e-04) (**Figure 5a**). Several genes in the fatty acid degradation (FAD) pathway displayed significant down-regulation in MM as compared to IN samples (*q* <0.1) (**Figure 5b**). Among the top 10 down-regulated genes in the FAD pathway, 5 out of 10 genes also showed significant down-regulation in the two metastatic cohorts (*P*<0.05, **Figure 5c**). *CYP4A11*, an enzyme of the cytochrome P450 family involved in fatty acid omega oxidation, displayed very strong down-regulation (>4-fold) across both, MM and SM, metastatic groups. Strikingly, *CYP4A11* is encoded at 1p33, displaying frequent copy number loss and concomitant gene expression down-regulation in MM tumors (**Figure 2e**). Fatty acids (FAs), a source of energy and essential components of bio-membranes, are degraded through three distinct pathways: α-, β- and ω- oxidation. We observed down-regulation of gene expression along all three different routes of fatty acid oxidation in MM as compared to IN tumors (**Figure 5d** and methods). Overall, these results suggest that down-regulation of FAD, potentially leading to the accumulation of FAs and lipid droplets (‘clear cell’ phenotype) inside the cancer cells, could be a feature of cancer cells at risk of metastatic progression.

**Figure 5:**
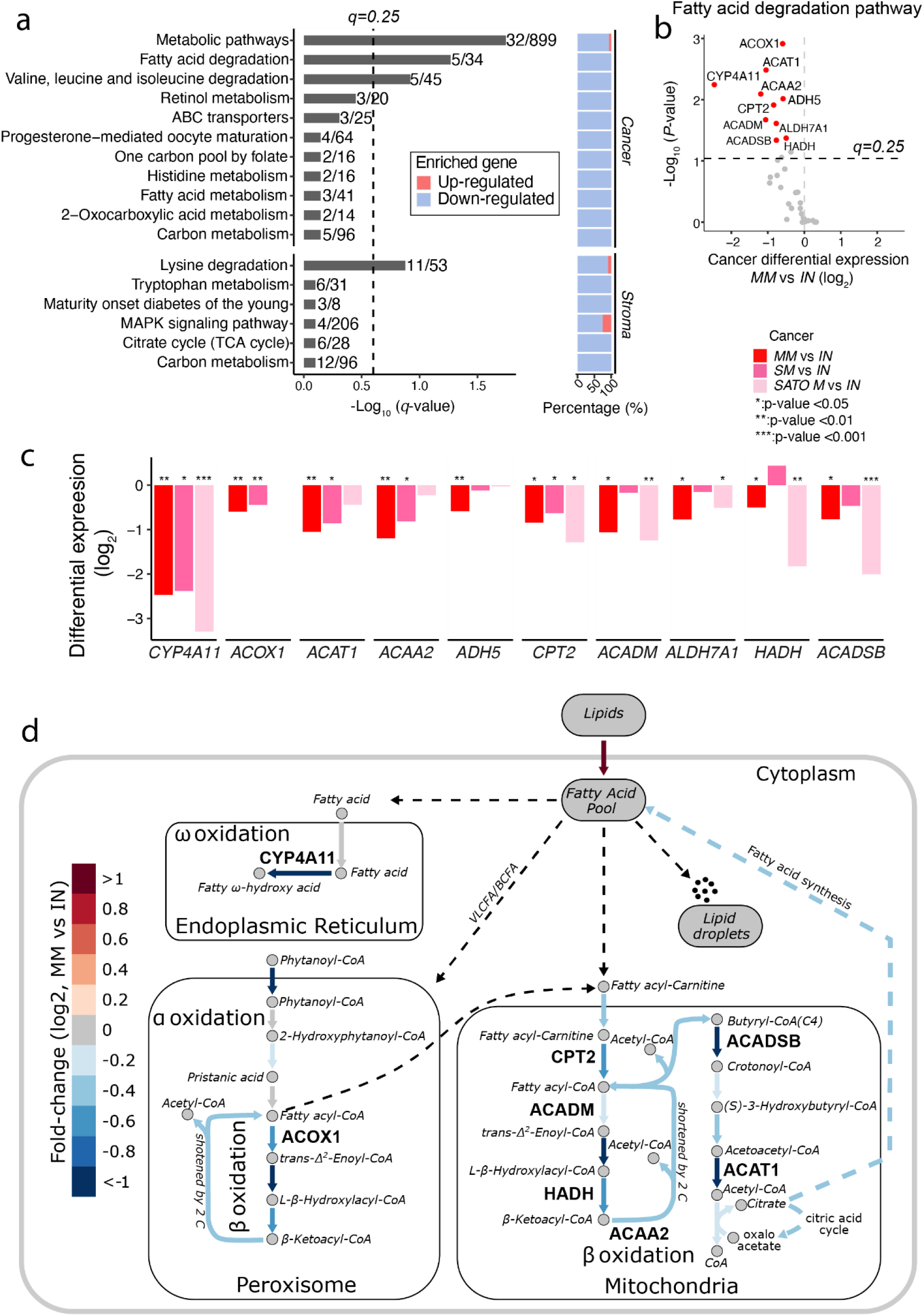
Fatty acid metabolism associated with risk of metachronous metastasis. **a**) Pathway enrichment of cancer and stromal differentially expressed genes in MM vs IN cohorts (Figure 4b, *P*<0.05). The proportion of up and down-regulated genes is indicated for each pathway. **b**) Inferred cancer-cell expression differences (MM vs IN) of genes involved in the fatty acid degradation (FAD) pathway. **c**) Differential expression of significant FAD genes (q<0.1) across groups and external SATO- M cohort. **d**) Map of FAD metabolic reactions. Metabolites and reactions are indicated with circles and arrows, respectively. Reactions are color-coded according to cancer-cell expression changes (MM vs. IN) of genes/enzymes in the given reaction. Genes highlighted in panel c are labelled on the map.

### Predicting risk of metachronous metastasis using a 5-gene signature

Pre-metastatic molecular features associated with the risk of developing metachronous metastasis could be incorporated into a model to predict risk of metastatic relapse. To develop such a model, we performed a Cox regression analysis on all putative genomic and molecular biomarker features identified in our study. Twenty-two features were individually predictive of metachronous metastasis risk (Benjamini-Hochberg*, q*<0.05, **Figure 6a**). We selected the top five predictive gene expression features (*FKBP15, SLC31A1, CPT2, PATJ* and *CALR*) to train a model (5G model) for predicting the risk of metachronous metastasis. We used a cross-validation approach to evaluate the accuracy of our model on unseen data, dividing the dataset into training (80%) and test sets (20%). We fitted multivariate Cox regression models using the signature on the training data, evaluating the error on the withheld test set (cross-validation). This process was repeated 100 times to estimate the average accuracy (AUC) of each model on the unseen test data. Previous studies have also proposed distinct gene signatures predictive of ccRCC disease recurrence (Brooks et al. 2014; Rini et al. 2015; Rappold et al. 2022) and mortality within 5 years of radical nephrectomy (Morgan et al. 2018). We compared the predictive accuracy of our two models with these existing signatures in stratifying MM from IN patients in our cohort. The 5G signature model demonstrated the best accuracy (73.7% AUC), with a specificity of 38.5% evaluated at 87.8% sensitivity. The existing Rini et al. (55.6% AUC, 16.24% specificity at 87.8% sensitivity), Morgan et al. (45.23% AUC, 10.67% specificity at 87.8% sensitivity), MSKI (56.32% AUC, 19.5% specificity at 87.8% sensitivity), clearcode34/ccB (60.92% AUC, 25.03% specificity at 87.8% sensitivity), and Alchahin et al. (61.25% AUC, 25.66% specificity at 87.8% sensitivity) signatures demonstrated overall lower accuracy in predicting metachronous metastasis risk (**Figure 6b)**. We further evaluated the association of the 5G model scores with disease-free survival. Splitting patients into tertiles based on the 5G score, the model demonstrated a significant difference in outcomes for patients with low and medium/high scores (**Figure 6c**, p=0.0088, log-rank test). Overall, these results suggest that the 5G model could be used to improve prediction of metastatic risk and stratify ccRCC patients into low and high-risk groups following curative surgery.

**Figure 6:**
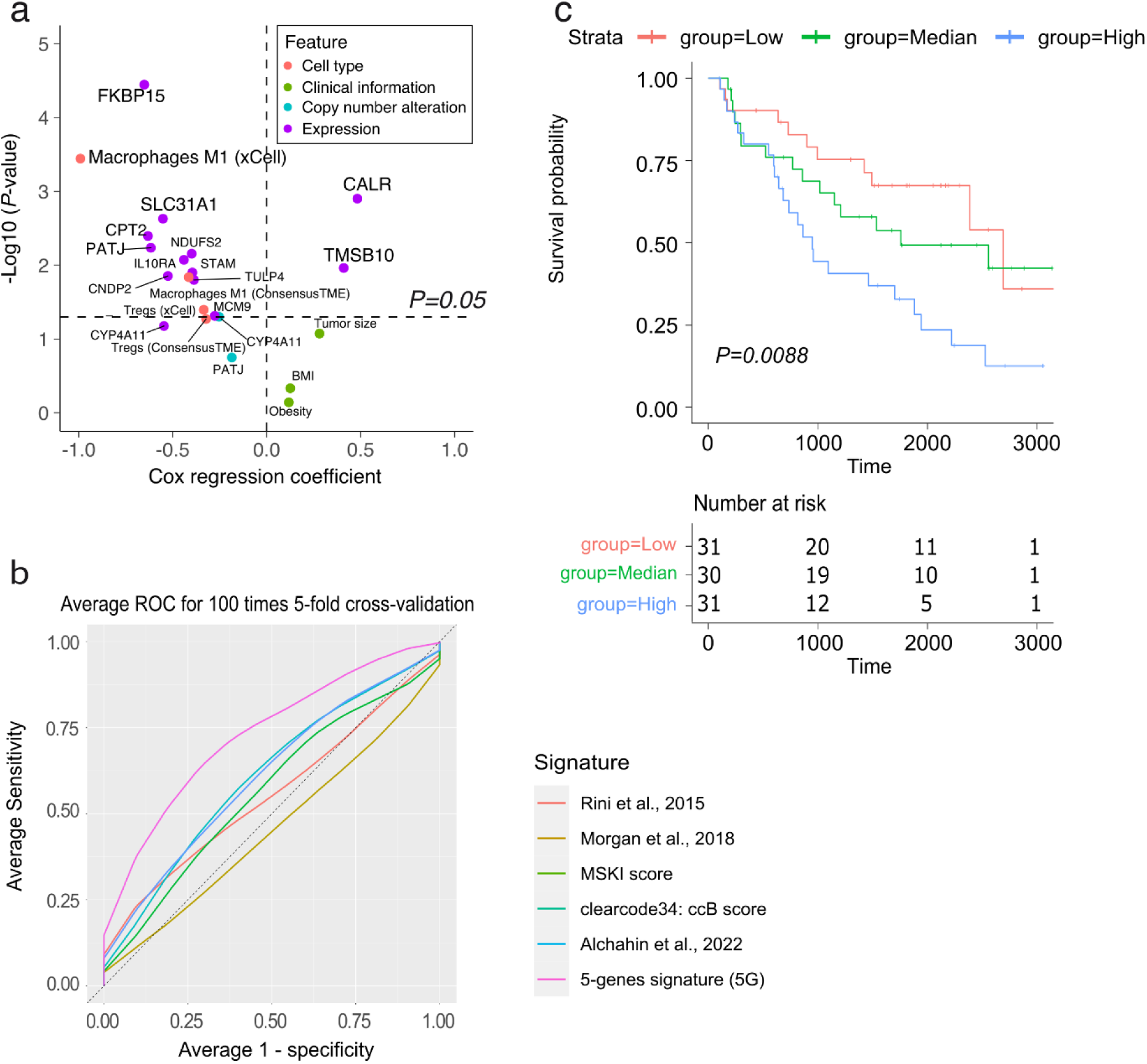
Genomic and molecular features predictive of metachronous metastasis risk. **a)** Cox regression analysis of individual candidate features associated with the risk of metachronous metastasis in this study. **b)** Average ROC for 100 times 5-fold cross-validation for prediction of metachronous metastasis risk; comparing the performance of the 5-gene (5G) signature including FKBP15, SLC31A1, CPT2, PATJ and CALR (purple) versus the ccRCC metastasis gene signatures previously published by Rini et al. 2015 (red), Morgan et al. 2018 (olive), MSKI score (green), clearcode34: ccB score (teal) and Alchahin et al 2022 (blue). **c)** Kaplan-Meier plot of disease-free survival (log-rank test) with the number of patients displayed for each group shown in the table below the plot.

To further validate the robustness of the 5G signature in an external dataset, we trained a cox model to predict overall survival using the 5G signature and evaluated its performance on the Sato dataset (Sato et al. 2013). Patients were divided into tertiles based on their 5G scores, yielding significant differences in outcomes for patients with predicted low, medium, and high-risk scores (**Supplementary** Figure 13, p=0.0012, log-rank test).

## Discussion

Our study aimed to identify cancer and stromal cell molecular features as putative biomarkers of localized ccRCC primary tumors at risk of metastatic progression. We investigated the genomic and transcriptomic profiles of 192 ccRCC primary tumors. In our study, these patients were analyzed using extended clinical follow-up beyond the standard TCGA clinical annotation. This allowed us to define subsets of high-confidence indolent tumors as well as tumors that at presentation appeared localized but later relapsed with metachronous metastasis (MM). We identified genomic aberrations as well as cancer and stroma cell gene expression signatures that characterize these primary tumors at risk of developing metastasis. Strikingly, these genomic and expression signatures converged at the 1p31-36 locus, displaying both frequent copy number loss and gene expression down-regulation in patients developing MM. Finally, these insights enabled us to develop a compact prognostic model to predict the risk of metastatic progression in ccRCC patients with localized disease.

Previous studies have reported altered frequencies of *BAP1* and *SETD2* driver mutations when comparing primary ccRCC tumors and metastatic tumors (Hakimi et al. 2013; Turajlic et al. 2018). Our study also identified an increased frequency of these mutations in metachronous metastatic tumors. However, this association was not statistically significant in our cohort, possibly due to a lower sample size and the restriction to stage III patients in this study. Instead, our analysis revealed that primary tumors display an increased probability of metastatic progression when they harbor specific genomic copy number aberrations. Specifically, we demonstrated that loss of chromosome 1p and gains of 2p, 2q, 19p and 19q were associated with the risk of metachronous metastasis. Consistent with previous studies of metastatic ccRCC lesions (Turajlic et al. 2018), we also found that 2q14.3 gain and 1p36.11 loss are more prevalent in tumors that later progress with metastasis. Moreover, our integrative analysis found that the 1p31-36 region (especially 1p33) also harbored genes, including *PATJ* and *CYP4A11*, that were significantly down-regulated in tumors that progressed with metastasis.

Using bulk tumor gene expression deconvolution, we observed significant down-regulation of *PATJ* expression in cancer cells of tumors that progressed with metachronous metastasis. *PATJ* encodes for a component of the Crumbs cell polarity complex and is involved in the formation of tight junctions as well as the apicobasal cell polarity (Shin et al. 2005, Pénalva and Mirouse 2012). Further analysis of factors involved in the regulation of tight junctions and cell polarity (Crumbs, Par, Scribble complexes) identified additional genes (*AMOLT2*, *CGN* and *TJP2*) with down-regulation in tumors that developed metastasis. Additionally, the top up-regulated gene in cancer cells of MM tumors was *TMSB10,* which has been linked to the regulation of cell polarity and metastatic invasion in breast cancer and ccRCC (Zhang et al. 2017, Pan et al. 2020). Consistent with existing knowledge on the role of these biological processes in metastatic seeding and progression (Martin-Belmonte and Perez-Moreno 2012, Gandalovičová et al. 2016, Salvador et al. 2016), our data suggest that primary ccRCC cancer cells with dysregulated epithelial cell polarity and tight junction formation exhibit an elevated risk of progressing with metastasis.

Our analysis also highlighted a link between metabolic reprogramming and the risk of metastatic progression. A hallmark of ccRCC is the accumulation of fatty acids (FAs) as lipid droplets inside cancer cells, the ‘clear-cell’ phenotype (Qiu et al. 2015). Fatty acids (FAs) are a source of energy and constitute essential components of bio-membranes. FAs are degraded through three distinct pathways including α-, β- and ω-oxidation. Here we report the down-regulation of gene expression along all three different pathways of fatty acid oxidation in tumors that later progressed with metastasis. In the ω- oxidation pathway, *CYP4A11* displayed very strong down-regulation (>4-fold) across all metastatic cohorts in our analysis. Intriguingly, *CYP4A11* is also encoded on the 1p33 chromosome locus displaying frequent copy number loss and gene expression down-regulation in tumors at risk of metastatic progression. We also observed strong down-regulation of Palmitoyltransferase-2 (*CPT2*), which is crucial for FA β-oxidation through the carnitine shuttle system into the mitochondrial matrix. FA β-oxidation has been demonstrated to be suppressed by hypoxia-inducible factors (HIFs) and is possibly essential for ccRCC tumorigenesis (Du et al. 2017). Intriguingly, our study indicates that tumors with reduced capacity for FA oxidation also exhibit a more aggressive phenotype with an elevated risk of metastatic progression. These findings should also be interpreted in the context that ccRCC tumors developing in an obesogenic environment may generally be more indolent (Hakimi et al. 2013).

We also identified a potential role for the tumor microenvironment (TME) in tumors at risk for metachronous metastasis. Using multiple immune cell deconvolution algorithms, we identified significant depletion of T-regulatory (Tregs) and M1 macrophage cell subsets in tumors at risk of metastatic progression. Consistent with these results, elevated Treg abundance in the TME has been observed in patients with favorable outcomes across diverse tumor types (Ward-Hartstonge and Kemp 2017, Givechian et al. 2018). Moreover, M1 macrophages are generally considered ‘good’ macrophages associated with increased anti-tumor inflammation (Aras and Zaidi 2017). Notably, a recent study suggested that intratumoral myeloid inflammation is associated with metastatic progression (Rappold et al. 2022). In line with our result, this study highlighted the enrichment of M1 macrophages in control mice without metastatic progression. In contrast, it has been suggested that M2 macrophages are associated with exhausted CD8 T-cells, advanced tumor stages, and metastatic progression (Braun et al. 2021, Rappold et al. 2022). Altogether, our results are concordant with these previous observations that metastatic progression of ccRCC is associated with a unique myeloid inflammatory state depleted from M1 macrophages.

Finally, we developed and evaluated a compact predictive model based on the molecular features associated with patient tumors developing metachronous metastasis. The 5G model, incorporating *FKBP15, SLC31A1, CPT2, PATJ,* and *CALR* expression, demonstrated improved accuracy in predicting risk of metastatic progression as compared to existing gene signatures proposed for the prediction of disease recurrence and mortality in ccRCC patients (Brooks et al. 2014, Rini et al. 2015, Morgan et al. 2018, Alchahin et al. 2022, Rappold et al. 2022). We validated the risk stratification performance in an external cohort (Sato et al. 2013); however, additional validation needs to be conducted to further evaluate the robustness of this model. The potential clinical utility for identifying patients at risk of developing metachronous metastasis must also be evaluated in combination with existing clinical risk models for ccRCC (Du et al. 2017).

In summary, we have identified convergent genomic and transcriptomic features associated with the risk of developing metachronous metastasis in ccRCC. Intriguingly, our study highlights chromosome 1p31-36 copy number loss, potentially driving dysregulation of both apicobasal cell polarity and fatty acid metabolism, as a key risk factor of metastatic seeding and progression in ccRCC. Our study also highlights a putative biomarker signature for the stratification of primary ccRCC tumors at risk of metastatic progression. Overall, these results provide new insights into the biology of metastatic ccRCC progression as well as potential new avenues for clinical management and risk stratification of early stage ccRCC patients.

## Methods

### Patient cohorts

We obtained data for 537 primary ccRCC tumor samples through The Cancer Genome Atlas (TCGA). Level 3 RNA-seq profiles (RNAseqV2 normalized RSEM) were downloaded from the UCSC Xena browser. We obtained Mutation Annotation files (MAF), level 3 Copy number variation and segmentation (CNA) from the Broad Institute GDAC Firehose website. Data URLs are provided in **Supplementary Table 1**.

We obtained curated clinical data with additional follow-up annotation of metastatic disease from the clinical TCGA (cTCGA) consortium. The median follow-up period was 23 months, with a maximum follow-up of 8 years. Cohorts were then stratified by median follow-up and patients’ metastatic status at the time of nephrectomy. Patients who developed metastasis within 100 days of surgery were categorized as synchronous metastatic cohort (SM, n=80). Patients with diagnosed metastatic disease after 100 days and up to 8 years were grouped as metachronous metastatic cohort (MM, n=44). Finally, patients that did not develop metastasis within their follow-up period were classified as indolent cohort (IN, n=68). The median follow-up for IN cohort was 51 months (**Supplementary Table 1**).

As a validation cohort, we used data from the Sato et al. (Sato et al. 2013). This dataset contained bulk tumor gene expression data (Agilent microarray), which was obtained from ArrayExpress. These samples (n=101) were stratified according to the TCGA cohort, yielding an SM (n= 32) and IN (n=69) cohort (**Supplementary** Figure 3). The median follow-up for the IN cohort was 51 months (**Supplementary Table 1**), equivalent to the TCGA IN cohort. We mapped the probe identifiers to HGNC gene symbols and applied arithmetic mean for single HGNC symbols with multiple mapped probe identifiers.

### Mutation and CNV analysis

Two-sided Fisher’s exact test was implemented to compare gene mutation frequencies of MM and SM tumors with IN tumors. Copy number alteration differences between cohorts were examined using a non-parametric two-tailed Wilcoxon rank-sum test. A two-tailed Wilcoxon rank-sum test was performed to compare bulk tumor expression of the genes located in 1p and 19q between MM and IN cohorts. Pearson correlation was used to examine correlations between CNV and bulk tumor expression in primary tumors, IN and MM samples.

### Tumor Purity Estimation

Tumor purity and intratumor heterogeneity were estimated using the approach described in Ghoshdastider et al. (Ghoshdastider et al. 2021). Estimates of tumor purity for TCGA cohort are based on a consensus of four different methods (PurBayes (Larson and Fridley 2013), ESTIMATE (Yoshihara et al. 2013), ASCAT (Van Loo et al. 2010), and AbsCN-seq (Bao et al. 2014)), using gene expression profiles, CNA segmentation, and SNV variant allele frequency (VAF) data of individuals. SATO purity estimates were inferred using PurBayes and ESTIMATE.

### Tumor gene expression deconvolution

We decomposed bulk tumor expression profiles to stroma and cancer cells expression using the TUMERIC model (Ghoshdastider et al. 2021). Bulk tumor expression values were log-transformed, log2(X+1), before deconvolution. Genes with median expression (FPKM)<2 were excluded from this analysis.

### Differential gene expression analysis

We used pair-wise class comparisons to identify genes differentially expressed between IN patients and the other two cohorts. To identify genes with altered expression restricted to a specific compartment (cancer vs. stroma), we developed a Differential Expression (DE) score. Log2 transformed expression values (*e*) inferred from TUMERIC for cancer (*c*) and stroma (*s*) were obtained. We then calculated the extent that a gene showed gene expression change (comparing groups 1 and 2) primarily in cancer or stroma compartment with the following DE score:

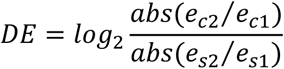

Here positive DE scores denote genes with higher expression change in the cancer compartment, and negative scores denote genes with higher differential expression in the stroma. A gene-wise two-tailed permutation test (non-parametric randomization test) was implemented to examine the statistical significance of the DE score. In this test, we estimated the *NULL* distribution of the DE statistic using random shuffles of the sample-to-group assignments. In each shuffle/permutation, cancer/stroma expression was estimated by TUMERIC and DE scores were computed. The *p*-values were evaluated based on n=50,000 permutations and computed using the fraction of *NULL* DE scores that are at least as extreme as the observed DE score for a given gene. The *p*-values were corrected for multiple testing using the Benjamini-Hochberg method. A similar method was performed for differential expression analysis in the SATO et al cohort.

### Cell Type Enrichment Analysis

Cell type deconvolution of cell types in the tumor microenvironment was performed using bulk tumor RNA-seq using the xCell R package (Aran et al. 2017), Consensus^TME^ (Jiménez-Sánchez et al. 2019), and Cibersort (Newman et al. 2015). xCell is a gene signature-based enrichment method that decomposes the expression data into 64 different cell types. Consensus^TME^ combines gene set-based methods and regression-based methods using multiple statistical tests to infer cell type enrichments. Cibersort is a deconvolution algorithm that estimates the abundances of 22 immune cell types using previous knowledge of immune cell expression signatures. To facilitate the analysis, Cibersort immune cells were grouped together as B cells (B Cells Naïve, B Cells Memory), T-Cell CD4 memory (T-Cell CD4 resting, T-Cell CD4 activated), T-Cell CD4 (T-Cell CD4 Naïve, T-Cell CD4 Memory), Natural Killer cells (NK Resting, NK Activated), macrophages (macrophage M0, macrophage M1, macrophage M2) as one group as well as separate groups, Dendritic cells (DC Activated, DC Resting), Mast cells (Mast cell resting, Mast cell activated), Monocytes, Plasma cells, Tgd cells and T regulatory cells (Tregs). Tumor samples with a Cibersort deconvolution performance p-value > 0.05 were excluded. Further analysis was performed on the shared cell types of the three methods with cell type fraction >10% in at least in 5 patients. To establish a cell type-wise differential analysis, we compared the median cell proportions of MM and IN patients. The significance level of these comparisons was estimated using a non-parametric Wilcoxon-rank-sum test. The *p*-values were adjusted to FDR *q*-values using the Benjamini-Hochberg method.

### Pathway Enrichment Analysis

To investigate differentially expressed genes in cancer and stroma cells, we used the KEGG pathway database (downloaded on September 2018, **Supplementary Table 1**) and pathway enrichment analysis (Ihaka and Gentleman 1996). The Bioconductor annotation package org.Hs.eg.db was used for mapping gene symbols to ENTREZ gene identifiers. The background gene list was defined as shared genes between KEGG pathways and the study datasets (n=4,505). To perform enrichment analysis in cancer and stroma discretely, two input gene lists were provided. These gene lists comprised the shared genes between the KEGG pathways and the significant genes in cancer (n=679) and stroma (n=720) gene expression analysis, as shown in Figure 6 (cancer and stroma *p*-values < 0.05). A one-sided Fisher’s exact test was applied to evaluate the significance level of pathway enrichment. *P*-values were adjusted for multiple testing using the Benjamini-Hochberg method.

### Tight junction and polarity complex analysis

To explore tight junction and polarity complexes, a gene list was derived from the tight junction pathway of the KEGG database. The 120 inferred genes were further curated and annotated as follows: Crumbs complex, Scribble complex, Par complex, Actin assembly, and others. Crumbs, Scribble, and Par complexes are the most well-known complexes responsible for cell polarity. Actin addresses the genes that participate in the structures and dynamics of the actin-based cytoskeleton. All other classifications are defined as ‘other regulators’.

### Metabolomic analysis

The Ensembl gene IDs in the RNA-seq data set were mapped to the Entrez IDs used in RECON3 using the BioMart R tool (Durinck et al. 2005, Durinck et al. 2009). RECON3 was manually checked and curated using information from literature, as previously described (Rohatgi et al. 2022). Furthermore, deconvoluted cancer and stromal cell expression for MM and IN tumors was integrated with the RECON3 based on the GPR information using the COBRA toolbox (Schellenberger et al. 2011). To estimate reaction expression from the deconvoluted cancer cell expression, we evaluated the GPR rule of each reaction while replacing the “OR” and “AND” operators with “max” and “min”, respectively (Colijn et al. 2009, Auslander et al. 2016, Rohatgi et al. 2022).

### Cox regression model predicting metachronous metastasis

To identify biomarkers associated with the risk of developing metachronous metastasis, we performed univariate Cox analyses on features (n=22) associated with metachronous metastasis in this study. We adjusted *p*-values using the Benjamini-Hochberg correction method. We selected the five most significant gene expressions (*FKBP15*, *SLC31A1*, *CPT2*, *PATJ* and *CALR*) as our signatures to predict metachronous metastasis. Using cross-validation, we then evaluated the predictive power and generalization performance of the identified 5G signature. We stratified samples into 5 folds, fitting multivariate Cox regression on 4 folds and evaluating the model performance on the remaining fold. We estimated matrices including the Akaike information criterion (AIC), the area under the curve (AUC), the concordance index (c-index), likelihood-ratio p-value, specificity and sensitivity to evaluate and compare the models. We repeated the analyses described above 100 times and calculated the mean value of all metrics. We compared the performance of the 5G signature for predicting metachronous metastasis with gene signatures provided by Brooks et al. 2014, Rini et al. 2015, Morgan et al. 2018, Alchahin et al. 2022 and Rappold et al. (Brooks et al. 2014, Rini et al. 2015, Morgan et al. 2018, Alchahin et al. 2022, Rappold et al. 2022).

### Log-rank survival analysis

We assessed whether the 5G score is associated with disease-free survival. Here we stratified patients into three groups based on their 5G scores. We performed a log-rank test using disease-free survival months and status for the three groups.

## Acknowledgments

MMN was supported by the A*STAR ARAP PhD scholarship. The results shown here are based upon data generated by the TCGA Research Network: https://www.cancer.gov/tcga. We are grateful to the patients who participated in this project.

## Author contributions

**MMN:** Conceptualization, Formal analysis, Methodology, Investigation, Visualization, Writing - Original Draft. **MP:** Formal analysis, Visualization, Investigation, Writing - Review & Editing. **NR:** Formal analysis, Visualization, Investigation, Writing - Review & Editing. **SK:** Investigation. **UG:** Formal analysis**. RGD:** Data curation, Resources. **RM:** Data curation, Resources. **AAH:** Conceptualization, Data curation, Resources, Investigation, Writing - Review & Editing. **AJS:** Conceptualization, Data curation, Methodology, Formal analysis, Investigation, Supervision, Project administration, Writing - Original Draft.

## Translational Relevance

Understanding the molecular features that determine the risk of metachronous metastases in clear cell renal cell carcinoma (ccRCC) is crucial for improving adjuvant treatment strategies and patient outcomes. We identified convergent genomic and transcriptomic alterations in chromosome 1p, implicating dysregulation of epithelial cell polarity and fatty acid metabolism as novel risk factors for metachronous metastases. Genomic analysis revealed a significantly higher frequency of copy number loss at chromosome 1p31-36 in primary tumors that later progressed with metastases. Integrative tumor transcriptome deconvolution further uncovered cancer and stromal cell molecular features associated with metastatic risk, independently implicating dysregulation of epithelial cell polarity and fatty acid metabolism genes at chromosome 1p (including *PATJ* and *CYP4A11*, at 1p31-33). Additionally, we developed, validated, and benchmarked a compact 5-feature predictive model (5G) that outperformed existing ccRCC gene signatures in predicting metachronous metastasis risk. Together, these findings offer new opportunities for drug development and improved patient risk management.

## References

1. Albiges, L., A. A. Hakimi, W. Xie, R. R. McKay, R. Simantov, X. Lin, J. L. Lee, B. I. Rini, S. Srinivas, G. A. Bjarnason, S. Ernst, L. A. Wood, U. N. Vaishamayan, S. Y. Rha, N. Agarwal, T. Yuasa, S. K. Pal, A. Bamias, E. C. Zabor, A. J. Skanderup, H. Furberg, A. P. Fay, G. de Velasco, M. A. Preston, K. M. Wilson, E. Cho, D. F. McDermott, S. Signoretti, D. Y. C. Heng and T. K. Choueiri (2016). "Body Mass Index and Metastatic Renal Cell Carcinoma: Clinical and Biological Correlations." J Clin Oncol 34(30): 3655–3663.

2. Alchahin, A. M., S. Mei, I. Tsea, T. Hirz, Y. Kfoury, D. Dahl, C.-L. Wu, A. O. Subtelny, S. Wu, D. T. Scadden, J. H. Shin, P. J. Saylor, D. B. Sykes, P. V. Kharchenko and N. Baryawno (2022). "A transcriptional metastatic signature predicts survival in clear cell renal cell carcinoma." Nature Communications 13(1): 5747.

3. Aran, D., Z. Hu and A. J. Butte (2017). "xCell: digitally portraying the tissue cellular heterogeneity landscape." Genome biology 18(1): 220–220.

4. Aras, S. and M. R. Zaidi (2017). "TAMeless traitors: macrophages in cancer progression and metastasis." British Journal of Cancer 117(11): 1583–1591.

5. Auslander, N., A. Wagner, M. Oberhardt and E. Ruppin (2016). "Data-Driven Metabolic Pathway Compositions Enhance Cancer Survival Prediction." PLoS computational biology 12(9): e1005125–e1005125.

6. Bao, L., M. Pu and K. Messer (2014). "AbsCN-seq: a statistical method to estimate tumor purity, ploidy and absolute copy numbers from next-generation sequencing data." Bioinformatics 30(8): 1056–1063.

7. Beroukhim, R., J.-P. Brunet, A. Di Napoli, K. D. Mertz, A. Seeley, M. M. Pires, D. Linhart, R. A. Worrell, H. Moch and M. A. Rubin (2009). "Patterns of gene expression and copy-number alterations in von-hippel lindau disease-associated and sporadic clear cell carcinoma of the kidney." Cancer research 69(11): 4674–4681.

8. Braun, D. A., K. Street, K. P. Burke, D. L. Cookmeyer, T. Denize, C. B. Pedersen, S. H. Gohil, N. Schindler, L. Pomerance, L. Hirsch, Z. Bakouny, Y. Hou, J. Forman, T. Huang, S. Li, A. Cui, D. B. Keskin, J. Steinharter, G. Bouchard, M. Sun, E. M. Pimenta, W. Xu, K. M. Mahoney, B. A. McGregor, M. S. Hirsch, S. L. Chang, K. J. Livak, D. F. McDermott, S. A. Shukla, L. R. Olsen, S. Signoretti, A. H. Sharpe, R. A. Irizarry, T. K. Choueiri and C. J. Wu (2021). "Progressive immune dysfunction with advancing disease stage in renal cell carcinoma." Cancer Cell 39(5): 632–648.e638.

9. Brooks, S. A., A. R. Brannon, J. S. Parker, J. C. Fisher, O. Sen, M. W. Kattan, A. A. Hakimi, J. J. Hsieh, T. K. Choueiri, P. Tamboli, J. K. Maranchie, P. Hinds, C. R. Miller, M. E. Nielsen and W. K. Rathmell (2014). "ClearCode34: A prognostic risk predictor for localized clear cell renal cell carcinoma." Eur Urol 66(1): 77–84.

10. Colijn, C., A. Brandes, J. Zucker, D. S. Lun, B. Weiner, M. R. Farhat, T.-Y. Cheng, D. B. Moody, M. Murray and J. E. Galagan (2009). "Interpreting expression data with metabolic flux models: predicting Mycobacterium tuberculosis mycolic acid production." PLoS computational biology 5(8): e1000489–e1000489.

11. Creighton, C. J., M. Morgan, P. H. Gunaratne, D. A. Wheeler, R. A. Gibbs, A. Gordon Robertson, A. Chu, R. Beroukhim, K. Cibulskis, S. Signoretti, F. Vandin Hsin-Ta Wu, B. J. Raphael, R. G. W. Verhaak, P. Tamboli, W. Torres-Garcia, R. Akbani, J. N. Weinstein, V. Reuter, J. J. Hsieh, A. Rose Brannon, A. Ari Hakimi, A. Jacobsen, G. Ciriello, B. Reva, C. J. Ricketts, W. Marston Linehan, J. M. Stuart, W. Kimryn Rathmell, H. Shen, P. W. Laird, D. Muzny, C. Davis, M. Morgan, L. Xi, K. Chang, N. Kakkar, L. R. Treviño, S. Benton, J. G. Reid, D. Morton, H. Doddapaneni, Y. Han, L. Lewis, H. Dinh, C. Kovar, Y. Zhu, J. Santibanez, M. Wang, W. Hale, D. Kalra, C. J. Creighton, D. A. Wheeler, R. A. Gibbs, G. Getz, K. Cibulskis, M. S. Lawrence, C. Sougnez, S. L. Carter, A. Sivachenko, L. Lichtenstein, C. Stewart, D. Voet, S. Fisher, S. B. Gabriel, E. Lander, R. Beroukhim, S. E. Schumacher, B. Tabak, G. Saksena, R. C. Onofrio, S. L. Carter, A. D. Cherniack, J. Gentry, K. Ardlie, C. Sougnez, G. Getz, S. B. Gabriel, M. Meyerson, A. Gordon Robertson, A. Chu, H.-J. E. Chun, A. J. Mungall, P. Sipahimalani, D. Stoll, A. Ally, M. Balasundaram, Y. S. N. Butterfield, R. Carlsen, C. Carter, E. Chuah, R. J. N. Coope, N. Dhalla, S. Gorski, R. Guin, C. Hirst, M. Hirst, R. A. Holt, C. Lebovitz, D. Lee, H. I. Li, M. Mayo, R. A. Moore, E. Pleasance, P. Plettner, J. E. Schein, A. Shafiei, J. R. Slobodan, A. Tam, N. Thiessen, R. J. Varhol, N. Wye, Y. Zhao, I. Birol, S. J. M. Jones, M. A. Marra, J. T. Auman, D. Tan, C. D. Jones, K. A. Hoadley, P. A. Mieczkowski, L. E. Mose, S. R. Jefferys, M. D. Topal, C. Liquori, Y. J. Turman, Y. Shi, S. Waring, E. Buda, J. Walsh, J. Wu, T. Bodenheimer, A. P. Hoyle, J. V. Simons, M. G. Soloway, S. Balu, J. S. Parker, D. Neil Hayes, C. M. Perou, R. Kucherlapati, P. Park, H. Shen, T. Triche Jr, D. J. Weisenberger, P. H. Lai, M. S. Bootwalla, D. T. Maglinte, S. Mahurkar, B. P. Berman, D. J. Van Den Berg, L. Cope, S. B. Baylin, P. W. Laird, C. J. Creighton, D. A. Wheeler, G. Getz, M. S. Noble, D. DiCara, H. Zhang, J. Cho, D. I. Heiman, N. Gehlenborg, D. Voet, W. Mallard, P. Lin, S. Frazer, P. Stojanov, Y. Liu, L. Zhou, J. Kim, M. S. Lawrence, L. Chin, F. Vandin, H.-T. Wu, B. J. Raphael, C. Benz, C. Yau, S. M. Reynolds, I. Shmulevich, R. G. W. Verhaak, W. Torres-Garcia, R. Vegesna, H. Kim, W. Zhang, D. Cogdell, E. Jonasch, Z. Ding, Y. Lu, R. Akbani, N. Zhang, A. K. Unruh, T. D. Casasent, C. Wakefield, D. Tsavachidou, L. Chin, G. B. Mills, J. N. Weinstein, A. Jacobsen, A. Rose Brannon, G. Ciriello, N. Schultz, A. Ari Hakimi, B. Reva, Y. Antipin, J. Gao, E. Cerami, B. Gross, B. Arman Aksoy, R. Sinha, N. Weinhold, S. Onur Sumer, B. S. Taylor, R. Shen, I. Ostrovnaya, J. J. Hsieh, M. F. Berger, M. Ladanyi, C. Sander, S. S. Fei, A. Stout, P. T. Spellman, D. L. Rubin, T. T. Liu, J. M. Stuart, S. Ng, E. O. Paull, D. Carlin, T. Goldstein, P. Waltman, K. Ellrott, J. Zhu, D. Haussler, P. H. Gunaratne, W. Xiao, C. Shelton, J. Gardner, R. Penny, M. Sherman, D. Mallery, S. Morris, J. Paulauskis, K. Burnett, T. Shelton, S. Signoretti, W. G. Kaelin, T. Choueiri, M. B. Atkins, R. Penny, K. Burnett, D. Mallery, E. Curley, S. Tickoo, V. Reuter, W. Kimryn Rathmell, L. Thorne, L. Boice, M. Huang, J. C. Fisher, W. Marston Linehan, C. D. Vocke, J. Peterson, R. Worrell, M. J. Merino, N. The Cancer Genome Atlas Research, M. Analysis working group: Baylor College of, B. C. C. Agency, I. Broad, Brigham, H. Women’s, U. Brown, M. D. A. C. C. The University of Texas, C. Memorial Sloan-Kettering Cancer, I. National Cancer, C. University of California Santa, C. H. University of North Carolina, C. University of Southern, M. Genome sequencing centres: Baylor College of, I. Genome characterization centres: Broad, S. Harvard Medical, C. University of Southern, U. Johns Hopkins, M. Genome data analysis: Baylor College of, A. Buck Institute for Research on, B. Institute for Systems, H. Oregon, U. Science, U. Stanford, H. University of, C. Biospecimen core resource: International Genomics, B. Tissue source sites, H. Women’s, I. Dana-Farber Cancer, U. Georgetown, C. International Genomics and H. University of North Carolina at Chapel (2013). "Comprehensive molecular characterization of clear cell renal cell carcinoma." Nature 499(7456): 43–49.

12. Du, C., F. Cai, M. A. Zidan, W. Ma, S. H. Lee and W. D. Lu (2017). "Reservoir computing using dynamic memristors for temporal information processing." Nature Communications 8(1): 2204.

13. Dudani, S., M.-F. Savard and D. Y. Heng (2019). "An update on predictive biomarkers in metastatic renal cell carcinoma." European urology focus.

14. Durinck, S., Y. Moreau, A. Kasprzyk, S. Davis, B. De Moor, A. Brazma and W. Huber (2005). "BioMart and Bioconductor: a powerful link between biological databases and microarray data analysis." Bioinformatics 21(16): 3439–3440.

15. Durinck, S., P. T. Spellman, E. Birney and W. Huber (2009). "Mapping identifiers for the integration of genomic datasets with the R/Bioconductor package biomaRt." Nature Protocols 4(8): 1184–1191.

16. Gandalovičová, A., T. Vomastek, D. Rosel and J. Brábek (2016). "Cell polarity signaling in the plasticity of cancer cell invasiveness." Oncotarget 7(18): 25022–25049.

17. Ghoshdastider, U., N. Rohatgi, M. Mojtabavi Naeini, P. Baruah, E. Revkov, Y. A. Guo, S. Rizzetto, A. M. L. Wong, S. Solai, T. T. Nguyen, J. P. S. Yeong, J. Iqbal, P. H. Tan, B. Chowbay, R. Dasgupta and A. J. Skanderup (2021). "Pan-Cancer Analysis of Ligand–Receptor Cross-talk in the Tumor Microenvironment." Cancer Research 81(7): 1802–1812.

18. Givechian, K. B., K. Wnuk, C. Garner, S. Benz, H. Garban, S. Rabizadeh, K. Niazi and P. Soon-Shiong (2018). "Identification of an immune gene expression signature associated with favorable clinical features in Treg-enriched patient tumor samples." npj Genomic Medicine 3(1): 14.

19. Hakimi, A. A., E. Reznik, C.-H. Lee, C. J. Creighton, A. R. Brannon, A. Luna, B. A. Aksoy, E. M. Liu, R. Shen, W. Lee, Y. Chen, S. M. Stirdivant, P. Russo, Y.-B. Chen, S. K. Tickoo, V. E. Reuter, E. H. Cheng, C. Sander and J. J. Hsieh (2016). "An Integrated Metabolic Atlas of Clear Cell Renal Cell Carcinoma." Cancer Cell 29(1): 104–116.

20. Hakimi, A. A. and M. H. Voss (2018). "Genomic Classifiers in Renal Cell Carcinoma." European urology 73(5): 770.

21. Ihaka, R. and R. Gentleman (1996). "R: a language for data analysis and graphics." Journal of computational and graphical statistics 5(3): 299–314.

22. Jiménez-Sánchez, A., O. Cast and M. L. Miller (2019). "Comprehensive Benchmarking and Integration of Tumor Microenvironment Cell Estimation Methods." Cancer Research 79(24): 6238–6246.

23. Larson, N. B. and B. L. Fridley (2013). "PurBayes: estimating tumor cellularity and subclonality in next-generation sequencing data." Bioinformatics 29(15): 1888–1889.

24. Martin-Belmonte, F. and M. Perez-Moreno (2012). "Epithelial cell polarity, stem cells and cancer." Nature Reviews Cancer 12(1): 23–38.

25. Morgan, T. M., R. Mehra, P. Tiemeny, J. S. Wolf, S. Wu, Z. Sangale, M. Brawer, S. Stone, C.-L. Wu and A. S. Feldman (2018). "A multigene signature based on cell cycle proliferation improves prediction of mortality within 5 yr of radical nephrectomy for renal cell carcinoma." European urology 73(5): 763–769.

26. Newman, A. M., C. L. Liu, M. R. Green, A. J. Gentles, W. Feng, Y. Xu, C. D. Hoang, M. Diehn and A. A. Alizadeh (2015). "Robust enumeration of cell subsets from tissue expression profiles." Nature Methods 12(5): 453–457.

27. Pan, Q., G. Cheng, Y. Liu, T. Xu, H. Zhang and B. Li (2020). "TMSB10 acts as a biomarker and promotes progression of clear cell renal cell carcinoma." International journal of oncology 56(5): 1101–1114.

28. Pénalva, C. and V. Mirouse (2012). "Tissue-specific function of Patj in regulating the Crumbs complex and epithelial polarity." Development 139(24): 4549–4554.

29. Qiu, B., D. Ackerman, D. J. Sanchez, B. Li, J. D. Ochocki, A. Grazioli, E. Bobrovnikova-Marjon, J. A. Diehl, B. Keith and M. C. Simon (2015). "HIF2α-dependent lipid storage promotes endoplasmic reticulum homeostasis in clear-cell renal cell carcinoma." Cancer discovery 5(6): 652–667.

30. Quail, D. F. and J. A. Joyce (2013). "Microenvironmental regulation of tumor progression and metastasis." Nature Medicine 19(11): 1423–1437.

31. Rappold, P. M., L. Vuong, J. Leibold, N. H. Chakiryan, M. Curry, F. Kuo, E. Sabio, H. Jiang, B. G. Nixon and M. Liu (2022). "A Targetable Myeloid Inflammatory State Governs Disease Recurrence in Clear-Cell Renal Cell Carcinoma." Cancer Discovery 12(10): 2308–2329.

32. Rini, B., A. Goddard, D. Knezevic, T. Maddala, M. Zhou, H. Aydin, S. Campbell, P. Elson, S. Koscielny and M. Lopatin (2015). "A 16-gene assay to predict recurrence after surgery in localised renal cell carcinoma: development and validation studies." The lancet oncology 16(6): 676–685.

33. Rohatgi, N., U. Ghoshdastider, P. Baruah, T. Kulshrestha and A. J. Skanderup (2022). "A pan-cancer metabolic atlas of the tumor microenvironment." Cell Reports 39(6): 110800.

34. Salvador, E., M. Burek and C. Y. Förster (2016). "Tight Junctions and the Tumor Microenvironment." Current Pathobiology Reports 4(3): 135–145.

35. Sato, Y., T. Yoshizato, Y. Shiraishi, S. Maekawa, Y. Okuno, T. Kamura, T. Shimamura, A. Sato-Otsubo, G. Nagae and H. Suzuki (2013). "Integrated molecular analysis of clear-cell renal cell carcinoma." Nature genetics 45(8): 860.

36. Scelo, G., Y. Riazalhosseini, L. Greger, L. Letourneau, M. Gonzàlez-Porta, M. B. Wozniak, M. Bourgey, P. Harnden, L. Egevad, S. M. Jackson, M. Karimzadeh, M. Arseneault, P. Lepage, A. How-Kit, A. Daunay, V. Renault, H. Blanché, E. Tubacher, J. Sehmoun, J. Viksna, E. Celms, M. Opmanis, A. Zarins, N. S. Vasudev, M. Seywright, B. Abedi-Ardekani, C. Carreira, P. J. Selby, J. J. Cartledge, G. Byrnes, J. Zavadil, J. Su, I. Holcatova, A. Brisuda, D. Zaridze, A. Moukeria, L. Foretova, M. Navratilova, D. Mates, V. Jinga, A. Artemov, A. Nedoluzhko, A. Mazur, S. Rastorguev, E. Boulygina, S. Heath, M. Gut, M.-T. Bihoreau, D. Lechner, M. Foglio, I. G. Gut, K. Skryabin, E. Prokhortchouk, A. Cambon-Thomsen, J. Rung, G. Bourque, P. Brennan, J. Tost, R. E. Banks, A. Brazma and G. M. Lathrop (2014). "Variation in genomic landscape of clear cell renal cell carcinoma across Europe." Nature Communications 5(1): 5135.

37. Schellenberger, J., R. Que, R. M. T. Fleming, I. Thiele, J. D. Orth, A. M. Feist, D. C. Zielinski, A. Bordbar, N. E. Lewis, S. Rahmanian, J. Kang, D. R. Hyduke and B. Ø. Palsson (2011). "Quantitative prediction of cellular metabolism with constraint-based models: the COBRA Toolbox v2.0." Nature protocols 6(9): 1290–1307.

38. Şenbabaoğlu, Y., R. S. Gejman, A. G. Winer, M. Liu, E. M. Van Allen, G. de Velasco, D. Miao, I. Ostrovnaya, E. Drill, A. Luna, N. Weinhold, W. Lee, B. J. Manley, D. N. Khalil, S. D. Kaffenberger, Y. Chen, L. Danilova, M. H. Voss, J. A. Coleman, P. Russo, V. E. Reuter, T. A. Chan, E. H. Cheng, D. A. Scheinberg, M. O. Li, T. K. Choueiri, J. J. Hsieh, C. Sander and A. A. Hakimi (2016). "Tumor immune microenvironment characterization in clear cell renal cell carcinoma identifies prognostic and immunotherapeutically relevant messenger RNA signatures." Genome Biology 17(1): 231.

39. Shin, K., S. Straight and B. Margolis (2005). "PATJ regulates tight junction formation and polarity in mammalian epithelial cells." The Journal of cell biology 168(5): 705–711.

40. Thul, P. J., L. Åkesson, M. Wiking, D. Mahdessian, A. Geladaki, H. Ait Blal, T. Alm, A. Asplund, L. Björk, L. M. Breckels, A. Bäckström, F. Danielsson, L. Fagerberg, J. Fall, L. Gatto, C. Gnann, S. Hober, M. Hjelmare, F. Johansson, S. Lee, C. Lindskog, J. Mulder, C. M. Mulvey, P. Nilsson, P. Oksvold, J. Rockberg, R. Schutten, J. M. Schwenk, Å. Sivertsson, E. Sjöstedt, M. Skogs, C. Stadler, D. P. Sullivan, H. Tegel, C. Winsnes, C. Zhang, M. Zwahlen, A. Mardinoglu, F. Pontén, K. von Feilitzen, K. S. Lilley, M. Uhlén and E. Lundberg (2017). "A subcellular map of the human proteome." Science 356(6340).

41. Tran, J. and M. C. Ornstein (2022). "Clinical Review on the Management of Metastatic Renal Cell Carcinoma." JCO Oncology Practice 18(3): 187–196.

42. Choueiri, T.K., Tomczak, P., Park, S.H., Venugopal, B., Ferguson, T., Symeonides, S.N., Hajek, J., Chang, Y.H., Lee, J.L., Sarwar, N. and Haas, N.B. (2024). "Overall survival with adjuvant pembrolizumab in renal-cell carcinoma." New England Journal of Medicine, 390(15): 1359–1371.

43. Turajlic, S., H. Xu, K. Litchfield, A. Rowan, T. Chambers, J. I. Lopez, D. Nicol, T. O’Brien, J. Larkin and S. Horswell (2018). "Tracking cancer evolution reveals constrained routes to metastases: TRACERx renal." Cell 173(3): 581–594. e512.

44. Van Loo, P., S. H. Nordgard, O. C. Lingjærde, H. G. Russnes, I. H. Rye, W. Sun, V. J. Weigman, P. Marynen, A. Zetterberg and B. Naume (2010). "Allele-specific copy number analysis of tumors." Proceedings of the National Academy of Sciences 107(39): 16910–16915.

45. Ward-Hartstonge, K. A. and R. A. Kemp (2017). "Regulatory T-cell heterogeneity and the cancer immune response." Clinical & translational immunology 6(9): e154–e154.

46. Wu, J., B.-Q. Shang, J.-Z. Shou and Y.-Y. Guan (2023). "A Novel Nomogram Predicting the Overall Survival of Patients with Metastatic Non-clear Cell Renal Cell Carcinoma: A Large Population-Based Investigation." Annals of Surgical Oncology.

47. Xiao, R., S. Shen, Y. Yu, Q. Pan, R. Kuang and H. Huang (2019). "TMSB10 promotes migration and invasion of cancer cells and is a novel prognostic marker for renal cell carcinoma." Int J Clin Exp Pathol 12(1): 305–312.

48. Yoshihara, K., M. Shahmoradgoli, E. Martínez, R. Vegesna, H. Kim, W. Torres-Garcia, V. Treviño, H. Shen, P. W. Laird and D. A. Levine (2013). "Inferring tumour purity and stromal and immune cell admixture from expression data." Nature communications 4: 2612.

49. Zhang, X., D. Ren, L. Guo, L. Wang, S. Wu, C. Lin, L. Ye, J. Zhu, J. Li, L. Song, H. Lin and Z. He (2017). "Thymosin beta 10 is a key regulator of tumorigenesis and metastasis and a novel serum marker in breast cancer." Breast Cancer Research 19(1): 15.

