## Supplementary figure 1-13 for "Convergent genomic and molecular features predict risk of metachronous metastasis in clear cell renal cell carcinoma"

Supplementary Figure 1: Overview of the follow-up time for indolent patients in SATO and TCGA cohorts.

Supplementary Figure 2: Overview of clinicopathological features in IN, MM and SM cohorts.

Supplementary Figure 3: Overview of Sato et al. cohort.

Supplementary Figure 4: Somatic mutations of FREM1 in MM cohort of TCGA.

Supplementary Figure 5: Copy number alterations and the correlations with expression at 1p31-36 chromosome region.

Supplementary Figure 6: Bulk tumor expression and copy number alterations.

Supplementary Figure 7: Cell type enrichment analysis.

Supplementary Figure 8: Bulk tumor expression of xCell Treg gene signatures.

Supplementary Figure 9: Bulk tumor expression of xCell M1 MPs gene signatures.

Supplementary Figure 10: Principal Component Analysis (PCA) using 3000 genes with the highest coefficient of variation.

Supplementary Figure 11: Deregulation of the cell polarity components.

Supplementary Figure 12: Normal kidney tissue single-cell RNA-seq data from the Human Protein Atlas (HPA).

Supplementary Figure 13: Kaplan-Meier plot of overall survival (log-rank test) in Sato dataset with the number of patients displayed for each group shown in the table below the plot.

Supplementary Table 1: Data URLs used in this study.

Supplementary Table 2: Differentially expressed genes in cancer cells of MM cohort versus IN cohort.

Supplementary Table 3: Differentially expressed genes in stromal cells of MM cohort versus IN cohort.

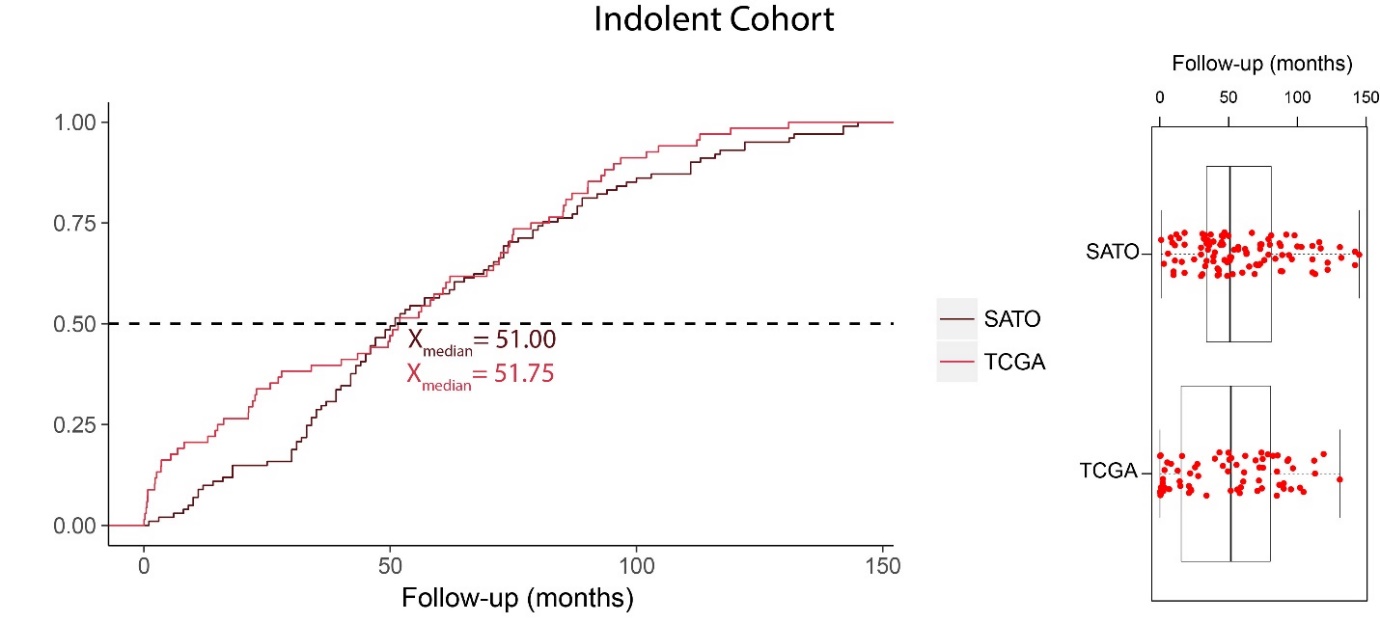

**Supplementary Figure 1**: **Overview of the follow-up time for indolent patients in SATO and TCGA cohorts.** Cumulative and box plots represent the median follow-up time of indolent patients at 51 months in SATO and 51.75 months in TCGA cohorts. Box plots showing the median values with the interquartile range (lower and upper hinge) and ±1.5-fold the interquartile range from the first and third quartile (lower and upper whiskers).

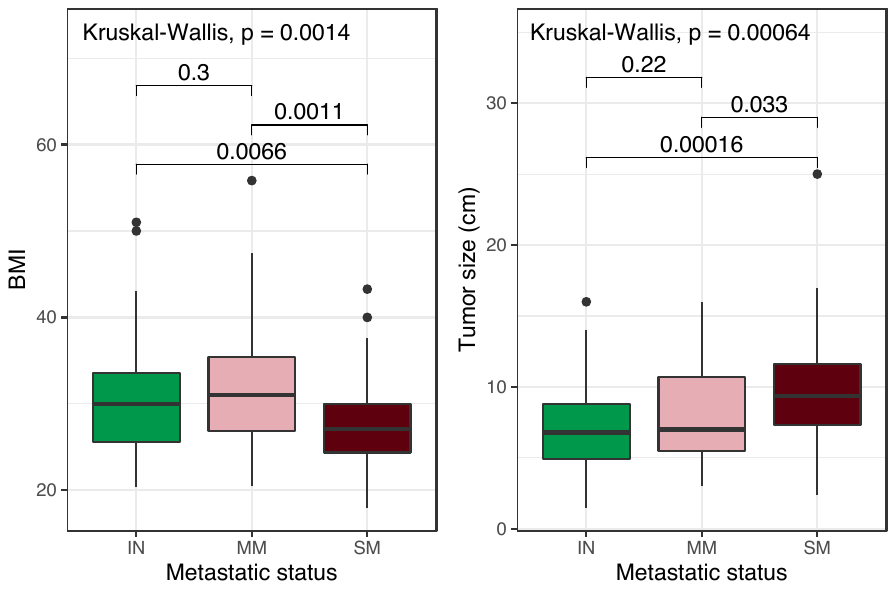

**Supplementary Figure 2**: **Overview of clinicopathological features in IN, MM and SM cohorts.** Box plots represent the distribution of BMI and tumor size across IN, MM and SM cohorts (Kruskal-Wallis for multiple group comparisons and Wilcoxon rank-sum test for pairwise mean comparisons). Box plots showing the median values with the interquartile range (lower and upper hinge) and ±1.5-fold the interquartile range from the first and third quartile (lower and upper whiskers).

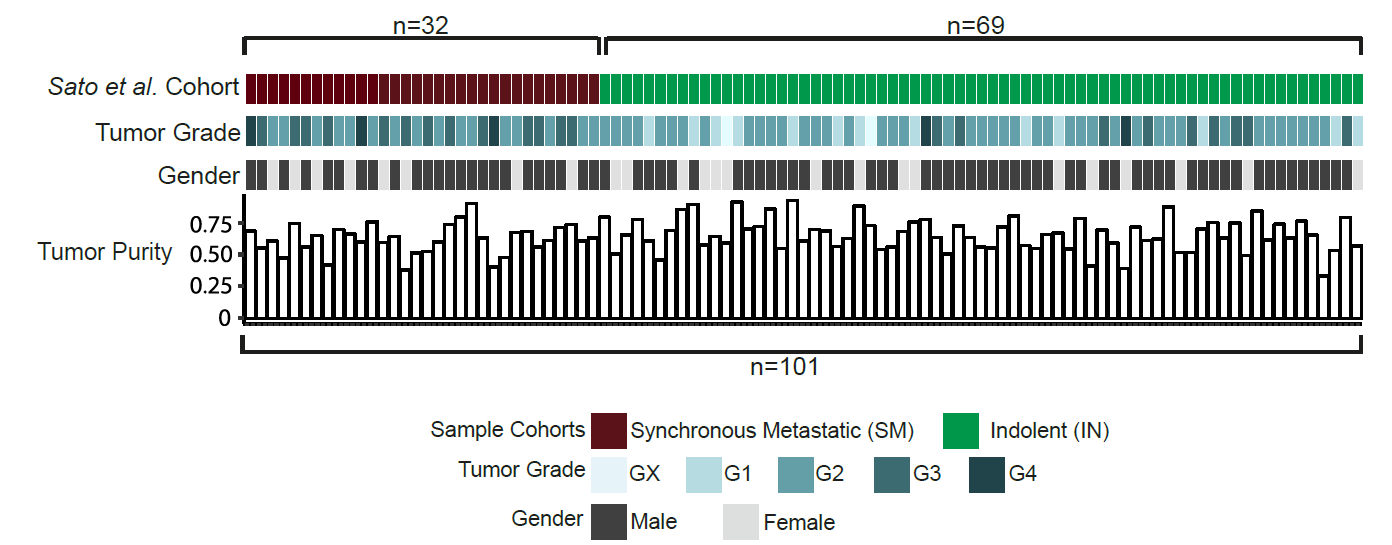

**Supplementary Figure 3**: **Overview of Sato et al. cohort (Sato et al., 2013)**. From top to bottom: sample cohorts, tumor grades, gender and estimated tumor purity. N, number.

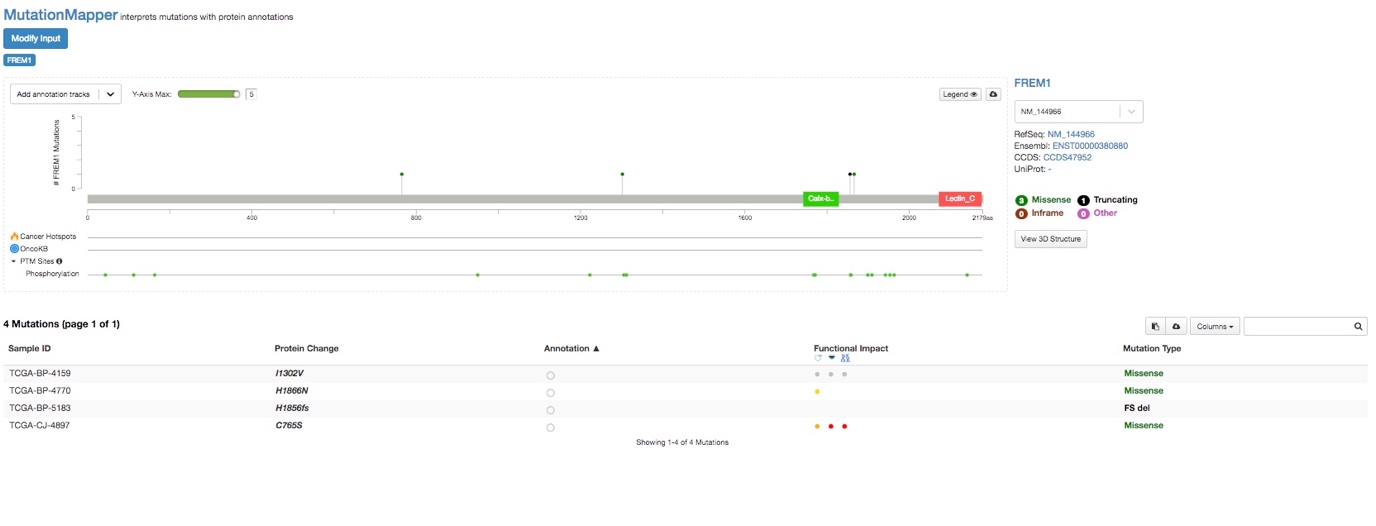

**Supplementary Figure 4**: **Somatic mutations of *FREM1* in MM cohort of TCGA.** Lollipop plot, mutation and protein annotations of *FREM1* somatic mutations in MM cohort of TCGA.

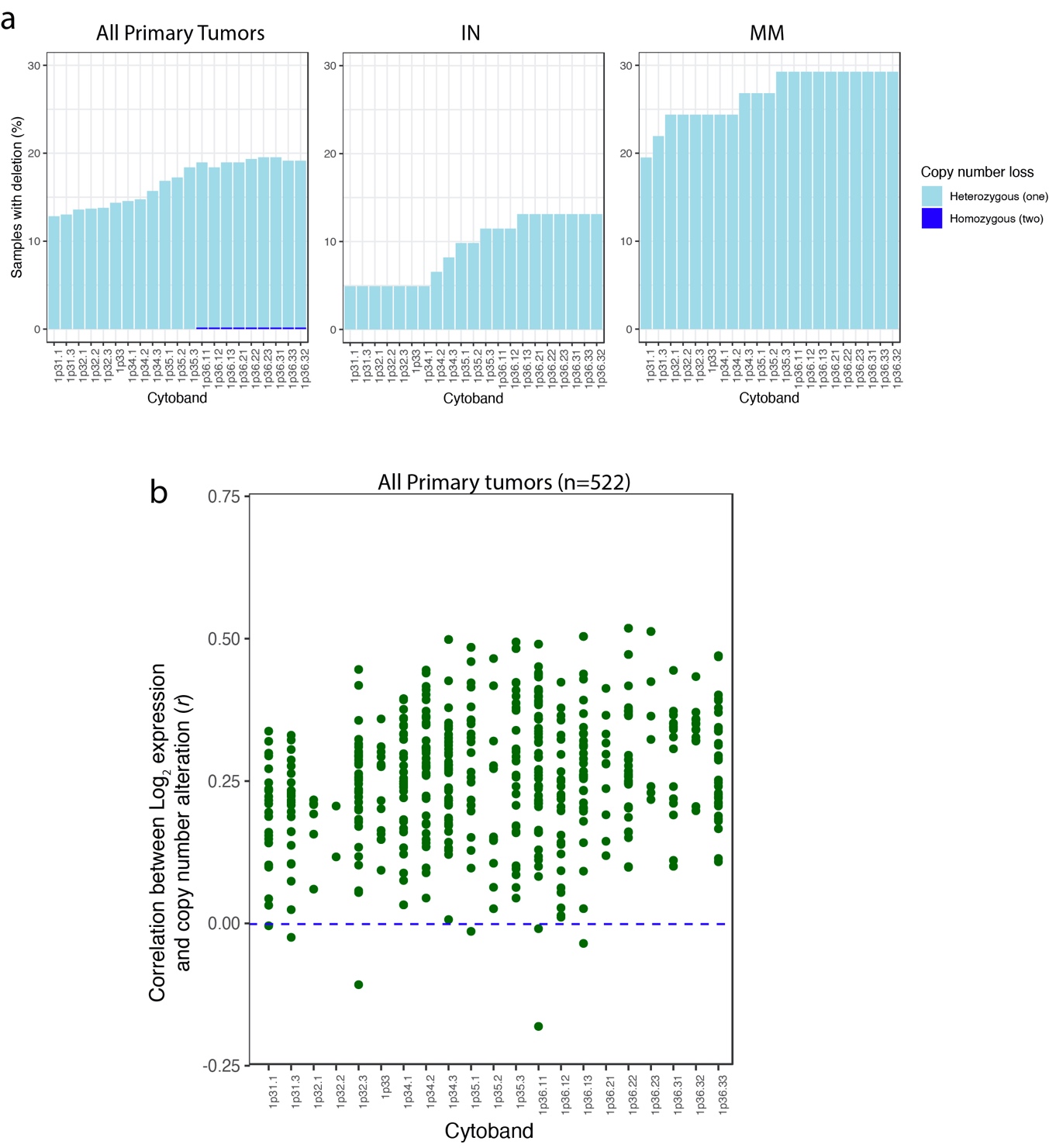

**Supplementary Figure 5**: **Copy number alterations and the correlations with expression at 1p31-36 chromosome region. a)** Frequency of samples with deletions in the 1p31-36 chromosome region. Left panel, all samples of renal clear cell carcinoma primary tumors (n=522); middle panel, IN cohort (n=61); right panel, MM cohort (n=41). One and two copy number losses are shown as light blue and dark blue, respectively. b) correlation of copy number alterations and expression values at 1p31-36 in all renal clear cell primary tumors of TCGA (n=522). N, number.

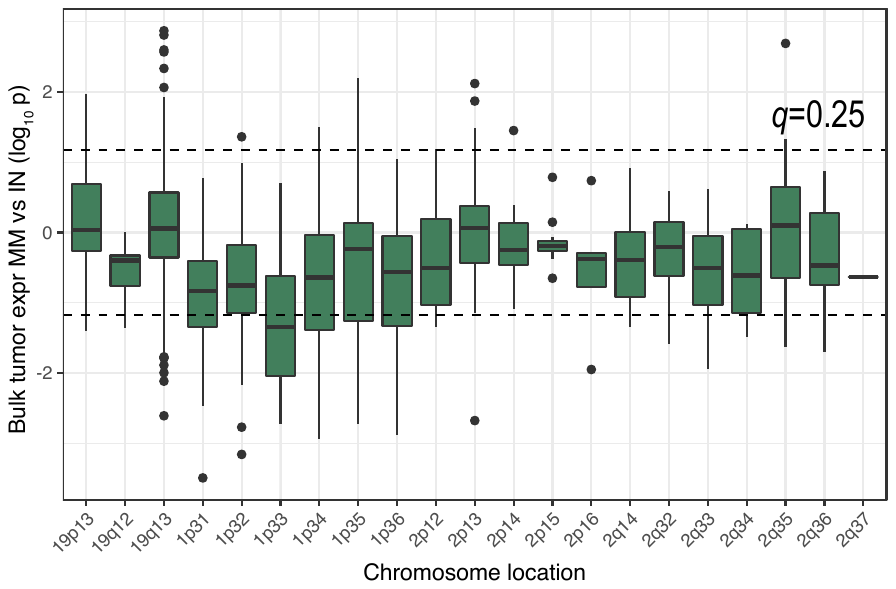

**Supplementary Figure 6**: **Bulk tumor expression and copy number alterations**. Differential bulk tumor expression in MM compared to IN tumor samples for genes located at 19p13-19q13, 1p31-36, 2p12-2p16, and 2q32-37 chromosome regions. The dashed line represents q=0.25 (Wilcoxon rank-sum test).

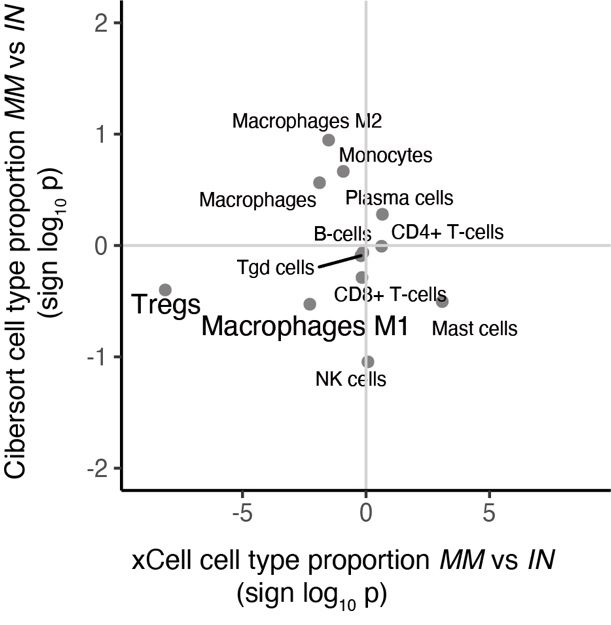

**Supplementary figure 7**: **Cell type enrichment analysis**. Differential analysis of cell type enrichment comparing MM and IN patients. Cell-type enrichment analysis was performed using xCell and Cibersort.

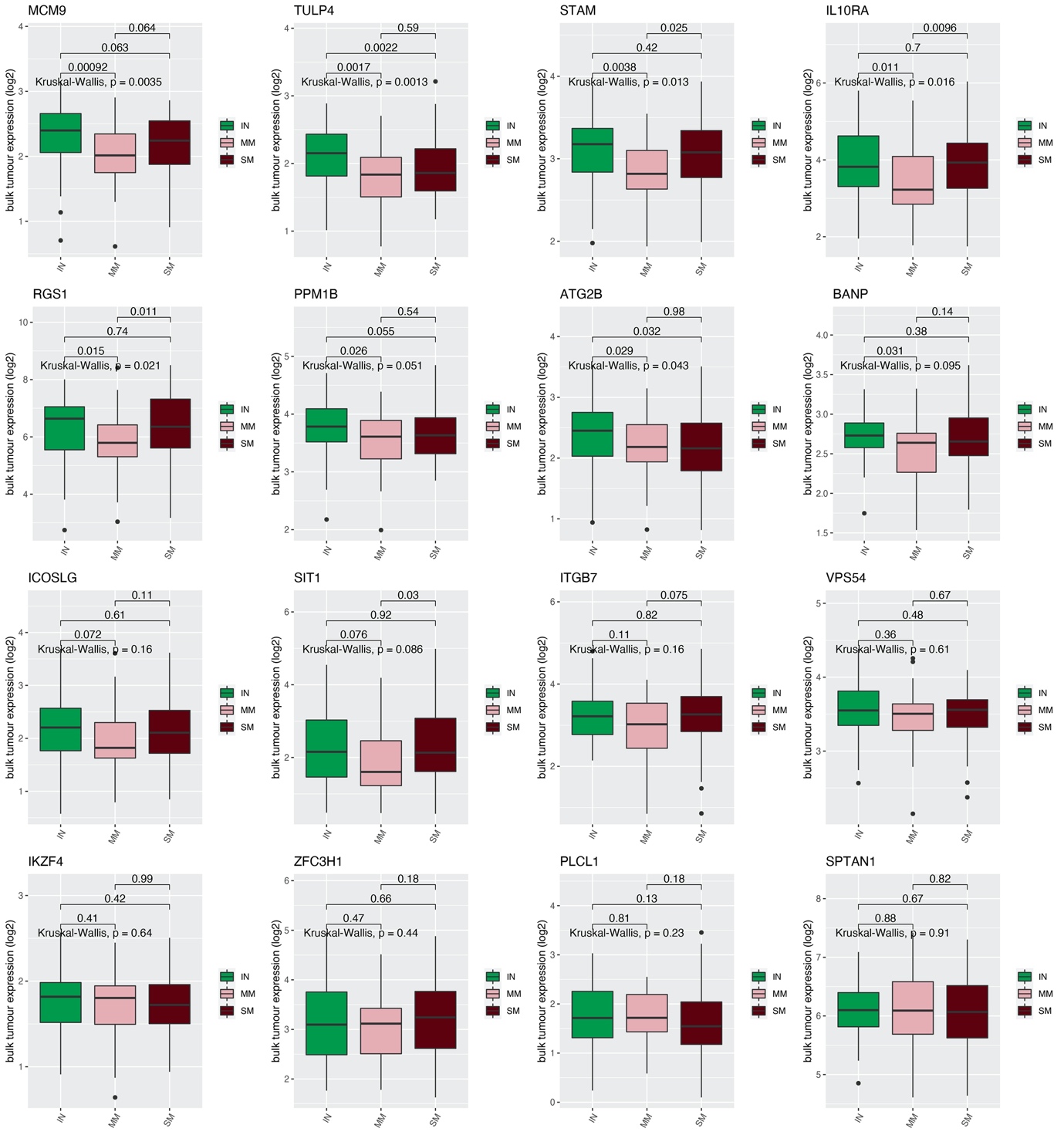

**Supplementary figure 8: Bulk tumor expression of xCell Treg gene signatures**. Box plots of bulk tumor expression for Treg gene markers in IN, MM and SM cohorts. Box plots showing the median values with the interquartile range (lower and upper hinge) and ±1.5-fold the interquartile range from the first and third quartile (lower and upper whiskers).

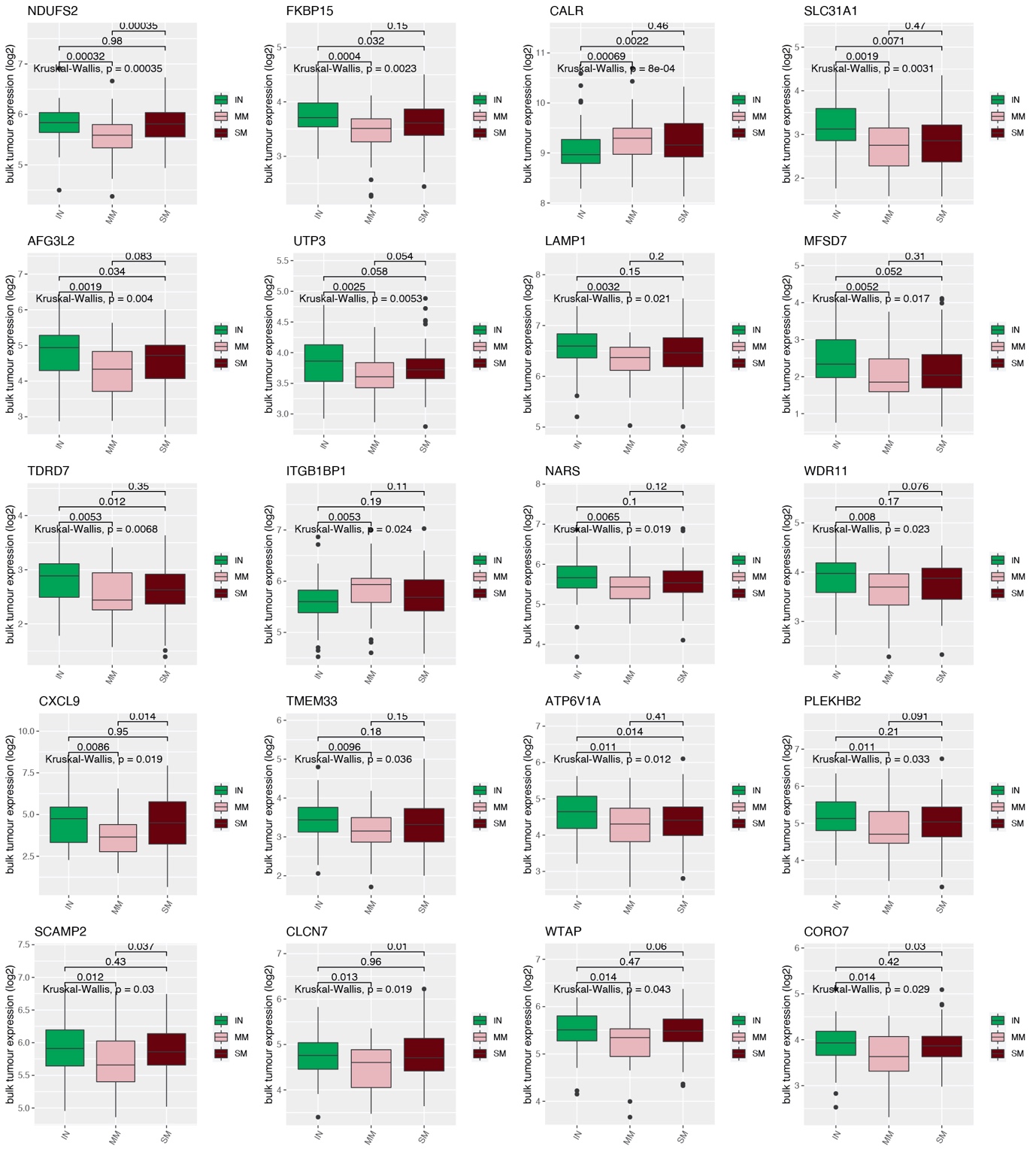

**Supplementary figure 9: Bulk tumor expression of xCell** **M1 MPs gene signatures**. Box plots of bulk tumor expression for the top 20 significant genes in IN, MM and SM cohorts. Box plots showing the median values with the interquartile range (lower and upper hinge) and ±1.5-fold the interquartile range from the first and third quartile (lower and upper whiskers).

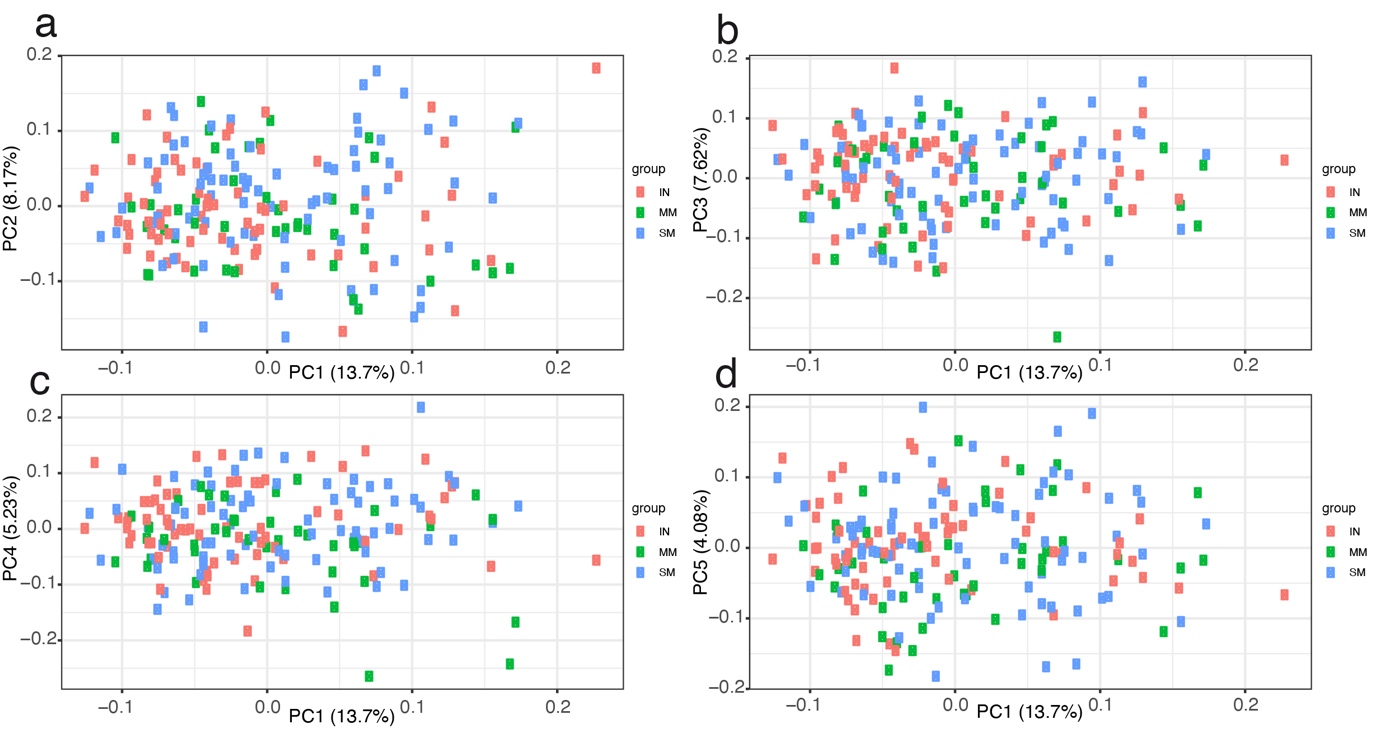

**Supplementary Figure 10**: **Principal Component Analysis (PCA) using 3000 genes with the highest coefficient of variation.** Displayed are the **a)** first (x-axis; PC1) and second (y-axis; PC2), **b)** first (x-axis; PC1) and third (y-axis; PC3), **c)** the first (x-axis; PC1) and fourth (y-axis; PC4), **d)** first (x-axis; PC1) and fifth (y-axis; PC5) Principal Components (PCs) of the genes for 189 samples. Color is representing IN, MM and SM groups.

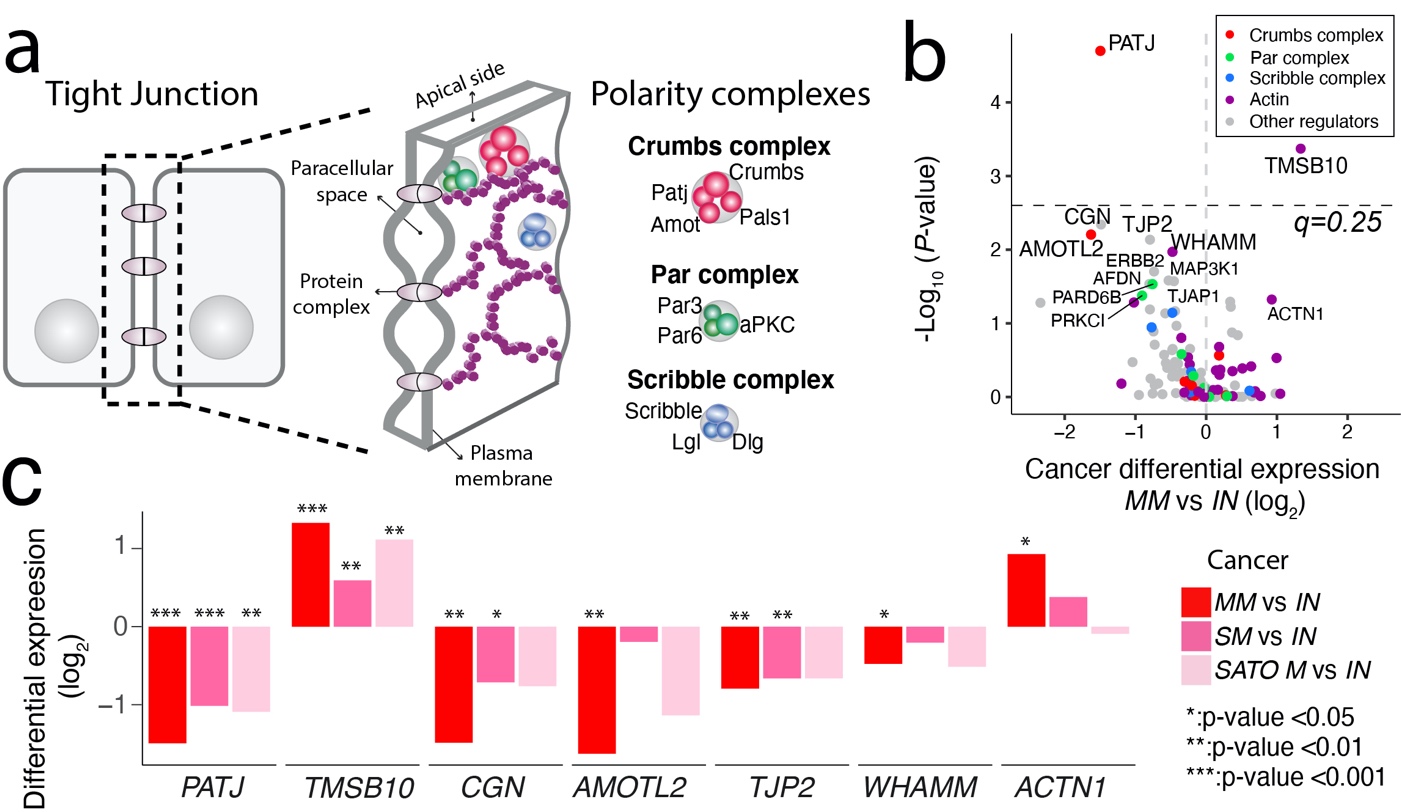

**Supplementary figure 11: Deregulation of the cell polarity components. a**) Schematic overview of tight junction and cell polarity components. Components of actin assembly (tight junction), Crumbs, Par and Scribble complexes are highlighted. **b**) Differential cancer compartment expression (MM vs IN) of 120 genes in the tight junction pathway (KEGG). Colors represent different polarity complexes shown in legend a). The dashed line represents *q*=0.25. **c**) Differential expression of the top significant tight junction genes in the MM, SM and SATO-M cohorts in comparison to IN tumors.

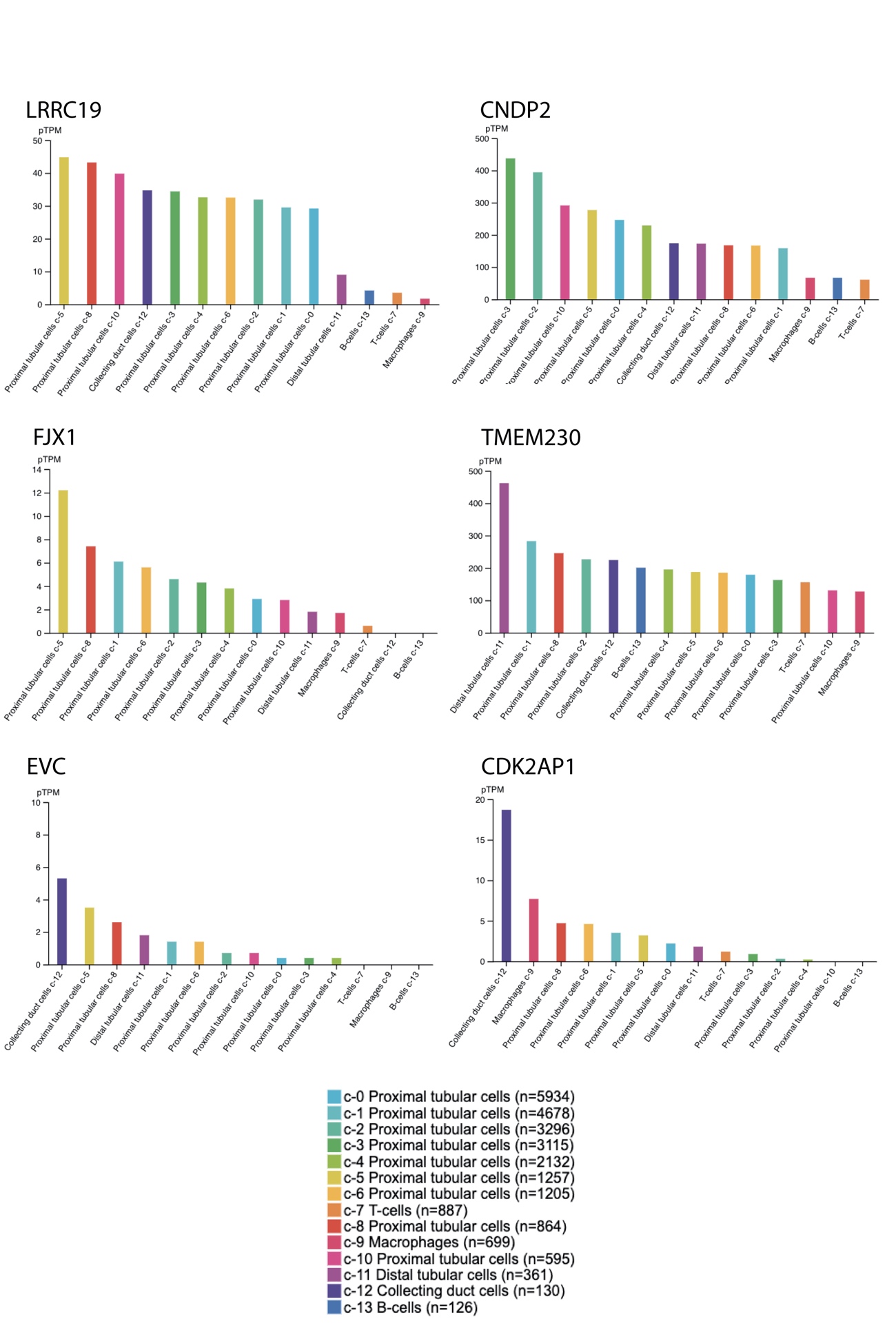

**Supplementary Figure 12**: **Normal kidney tissue single-cell RNA-seq data from the Human Protein Atlas (HPA).** Bar plots representing expression of LRRC19, CNDP2, FJX1, TMEM230, EVC and CDK2AP1 in normal tissue ordered from highest to lowest.

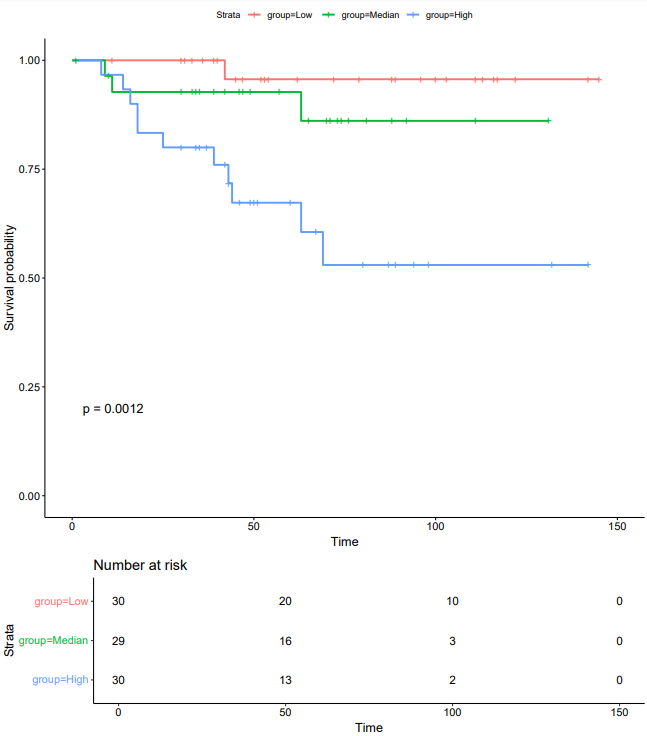

**Supplementary Figure 13**: **Kaplan-Meier plot of overall survival (log-rank test) in Sato dataset with the number of patients displayed for each group shown in the table below the plot.**

**References**

Sato, Y., Yoshizato, T., Shiraishi, Y., Maekawa, S., Okuno, Y., Kamura, T., . . . Suzuki, H. (2013). Integrated molecular analysis of clear-cell renal cell carcinoma. *Nature genetics, 45*(8), 860.
